# Proteomic Evaluation of the Progression from Acute Kidney Injury to Chronic Kidney Disease

**DOI:** 10.64898/2026.11.12.766273

**Authors:** Sunder Sims-Lucas, Joanna Bons, Katherine Pfister, Anne Silva-Barbosa, Rebecca Uhlean, Eric Goetzman, Birgit Schilling

**Affiliations:** University of Maryland Golisano Children’s Hospital, Baltimore, MD 21201, USA; Buck Institute for Research on Aging, Novato, CA 94945, USA; Department of Pediatrics, University of Pittsburgh Medical Center Children’s Hospital of Pittsburgh (UPMC CHP), University of Pittsburgh, Pittsburgh, PA 15224, USA

**Keywords:** acute kidney injury, chronic kidney disease, DIA-MS proteomics, AKI/CKD progression, peroxisomal dysfunction, mitochondrial dysfunction, cellular senescence, SASP

## Abstract

Acute kidney injury (AKI) and chronic kidney disease (CKD) are interconnected syndromes with substantial global burden, yet the molecular mechanisms driving the transition from acute injury to chronic fibrosis remain incompletely understood. We performed quantitative data-independent acquisition mass spectrometry (DIA-MS) proteomics on mouse models of AKI (ischemia-reperfusion injury) and CKD (unilateral ureteral obstruction), quantifying 5,985 protein groups. A conserved core injury response of 1,475 shared proteins (46%) was dominated by mitochondrial dysfunction, loss of solute transporters, and immune and extracellular matrix remodeling. AKI was marked by chromatin reorganization, DNA damage response, and acute inflammation, whereas CKD showed sustained fibrosis, complement activation, and coagulation. Peroxisomal proteins, including the β-oxidation enzymes Acox1 and Ehhadh, were consistently down-regulated in both AKI and CKD, with no peroxisomal protein up-regulated. Both models also displayed a senescence-associated secretory phenotype (SASP)-like signature, including Havcr1, Spp1, Igfbp7, Plau, and lysosomal cathepsins. We further identified 95 candidate progression markers, including Brat1 and Col3a1, 93 proteins with greater loss in CKD, 14 proteins changing in opposite directions, and seven disease-associated co-expression modules. These findings provide a proteomics resource for biomarker discovery and therapeutic targeting, including peroxisomal restoration and senolytic strategies, in AKI and CKD.

## Introduction

### Acute Kidney Injury and Chronic Kidney Disease: An Interconnected Clinical Syndrome

Acute kidney injury (AKI) represents a major clinical challenge, affecting 10–15% of hospitalized patients and more than 50% of those in intensive care units (1). The global burden of AKI continues to rise, with prevalence increasing from 15.5% in 2011 to 26.8% in 2021 (2). Historically, AKI and chronic kidney disease (CKD) were conceptualized as distinct clinical entities: AKI as an acute, reversible insult and CKD as a slowly progressive condition. However, compelling epidemiological and mechanistic evidence has fundamentally challenged this paradigm, establishing that AKI and CKD exist on a continuum with bidirectional interactions (3, 4). The recognition that AKI can initiate or accelerate CKD progression has transformed our understanding of kidney disease pathophysiology. Clinical studies have demonstrated that the severity, frequency, and duration of AKI episodes directly correlate with the risk of subsequent CKD development and progression to end-stage renal disease (5, 6). Even patients who appear to recover normal kidney function following an AKI episode remain at substantially elevated risk for long-term adverse outcomes. Glomerular filtration rate (GFR) decline following AKI has emerged as a critical marker, with decrements of greater than 25% from baseline strongly associated with increased mortality and CKD progression (7).

### Mechanisms of AKI-to-CKD Transition

The transition from AKI to CKD involves a complex interplay between maladaptive repair processes, persistent inflammation, and progressive fibrosis (8). Following acute injury, the balance between adaptive regeneration and maladaptive repair determines long-term outcomes. Tubular epithelial cells (TECs) serve as central orchestrators of this response, with damaged TECs secreting colony-stimulating factor 1 (CSF-1) that modulates macrophage recruitment and polarization (9). The cytokine milieu established following injury profoundly influences repair trajectory. Interleukin-1 (IL-1), IL-6, IL-10, and particularly transforming growth factor-β1 (TGF-β1) have been implicated in driving fibrotic responses through activation of resident fibroblasts and promotion of epithelial-to-mesenchymal transition (10, 11). TGF-β1, considered the master regulator of fibrosis, promotes myofibroblast differentiation, excessive extracellular matrix deposition, and perpetuation of the inflammatory cascade (12). Together, the progression to fibrosis is a hallmark of CKD.

Cell cycle dynamics following injury have emerged as critical determinants of repair outcomes. Tubular cells arrested in the G^2^/M phase of the cell cycle exhibit a profibrotic secretome characterized by elevated production of TGF-β1, connective tissue growth factor, and other fibrogenic mediators (13). This G^2^/M arrest is associated with activation of c-Jun N-terminal kinase (JNK) signaling, which further amplifies profibrotic cytokine production and inhibits completion of cell division (14).

Ferroptosis, a form of regulated cell death characterized by iron-dependent lipid peroxidation, has recently emerged as a significant contributor to AKI pathophysiology and the AKI-to-CKD transition (15). Unlike apoptosis, ferroptosis is driven by accumulation of lipid peroxides in cellular membranes, catalyzed by labile iron and insufficient antioxidant defenses (16). Inhibition of ferroptosis through iron chelation, lipid peroxide scavengers, or activation of the glutathione peroxidase 4 (GPX4) pathway has shown promise in ameliorating kidney injury in preclinical models (17, 18).

### Kidney Disease and Aging: A Growing but Underrecognized Burden

Global demographic shifts present unprecedented challenges for kidney disease management. By 2050, an estimated 1.5 billion people worldwide will be aged 65 years or older, representing a dramatic expansion of the population at highest risk for CKD (19). The relationship between aging and CKD is striking: prevalence increases from approximately 6.7% in individuals aged 40–59 years to 46.8% in those aged 70 years and older, with rates exceeding 50% in adults over 80 years (20, 21). Medicare data reveal that CKD prevalence approaches 10% in the elderly population compared to 1.5% in younger employed individuals (22).

Despite its magnitude, CKD remains significantly underrecognized compared to other age-related conditions such as neurodegeneration, osteoarthritis, and cancer. This disparity is particularly concerning given that CKD functions as a powerful risk multiplier, substantially increasing morbidity and mortality from diabetes, hypertension, cardiovascular disease, and stroke (23). Projections indicate that CKD could rise from the 16th leading cause of years of life lost globally in 2016 to the 5th leading cause by 2040, representing one of the largest projected increases among all disease categories (24).

Aging represents the single strongest predictor of CKD, reflecting both cumulative environmental exposures and intrinsic biological changes. The aging kidney undergoes characteristic structural and functional alterations, including nephron loss, glomerulosclerosis, tubulointerstitial fibrosis, and vascular rarefaction (25). These age-related changes create a reduced functional reserve that renders the elderly kidney particularly vulnerable to superimposed insults and impairs recovery following AKI.

Cellular senescence has emerged as a unifying mechanism linking kidney aging, AKI susceptibility, and CKD progression (26). Senescent cells, characterized by permanent cell cycle arrest and resistance to apoptosis, accumulate in aging tissues and adopt a senescence-associated secretory phenotype (SASP) that drives chronic low-grade inflammation (27). The SASP includes proinflammatory cytokines, chemokines, growth factors, and matrix metalloproteinases that collectively create a tissue microenvironment conducive to fibrosis and functional decline. Aging sensitizes tubular epithelial cells to G^2^/M arrest in response to stress and DNA damage, predisposing to senescence rather than successful repair (28). Studies using p16-reporter mice have revealed that senescent cells appear in proximal tubular segments within 1–3 days following ischemic injury (29). While early senescence may serve protective functions by limiting proliferation of damaged cells, persistence of senescent cells is strongly associated with poor recovery and CKD progression. Senolytic therapeutic strategies that selectively eliminate senescent cells have shown promise in preclinical models of kidney aging and disease (30).

### Proteomics in Kidney Disease Research

Proteomic workflows offer distinct advantages for kidney disease biomarker discovery and mechanistic investigation. Unlike antibody-based methods, mass spectrometric acquisitions enable untargeted, highly multiplexed protein identification without requiring a priori knowledge of target analytes (31). MS uniquely captures proteoform information, including post-translational modifications, truncations, and splice variants that may carry disease-specific signatures invisible to conventional immunoassays.

Current clinical diagnosis of AKI relies primarily on serum creatinine, a marker with well-documented limitations including delayed elevation that may not occur until approximately 50% of kidney function is lost (32, 33). Systematic proteomic investigations have identified promising plasma and urinary biomarker panels for AKI, including neutrophil gelatinase-associated lipocalin (NGAL), kidney injury molecule-1 (KIM-1), cystatin C, IL-18, α1-microglobulin, and fetuin-A (34–37). The CKD273 urinary peptide classifier, comprising 273 peptides identified by capillary electrophoresis-MS, provides superior early detection of CKD progression compared to conventional markers (38–40). More recently, a large-scale plasma proteomics study of 414 non-dialysis CKD patients (170 fast progressors and 244 stable patients) employed SWATH-MS and machine learning feature selection, and subsequently identified 25 plasma proteins as biomarkers of CKD progression, with enrichment of complement cascade and ubiquitin–proteasome pathways (41).

While biofluid proteomics has dominated kidney biomarker discovery, direct interrogation of kidney tissue provides mechanistic insights that cannot be obtained from plasma or urine. Integrated proteo-metabolomic studies of kidney tissue following ischemia-reperfusion injury have revealed coordinated changes in energy metabolism, including elevated acute phase response proteins, coagulation factors, and impaired mitochondrial ATP production alongside altered fatty acid metabolism (42). Our group has previously employed comprehensive data-independent acquisitions (DIA-MS) for deep proteomic profiling and succinylomics to characterize the proteomic landscape of AKI in murine models, establishing key signatures of mitochondrial dysfunction and tubular injury (43–46).

### Rationale and Study Objectives

Despite significant advances in understanding AKI-to-CKD transition mechanisms, substantial gaps remain in our ability to predict disease progression and identify therapeutic targets. Current biomarkers lack sufficient specificity and sensitivity for early detection of maladaptive repair, and the molecular heterogeneity underlying variable patient outcomes remains incompletely characterized. This study employs an in-depth quantitative DIA-MS proteomics approach to address these knowledge gaps using two complementary murine injury models (AKI and CKD). Our objectives are: (1) to identify novel protein biomarkers that distinguish adaptive from maladaptive kidney repair; (2) to elucidate molecular pathways driving the AKI-to-CKD transition; and (3) to identify potential therapeutic targets for preventing CKD progression following acute injury. By integrating differential expression analysis, pathway enrichment, subcellular localization mapping, and co-expression network analysis, we provide a comprehensive proteomics resource that bridges molecular discoveries to their cellular and functional contexts within the injured kidney.

## Results

### Proteomic profiling of acute and chronic kidney injury models

To comprehensively characterize the molecular signatures of acute versus chronic kidney injury, we performed quantitative DIA-MS proteomics on two well-established mouse models: ischemia-reperfusion injury (IRI) for AKI and unilateral ureteral obstruction (UUO) for CKD-associated fibrosis **(Figure 1A)**. As previously published (43–47), the IRI model involved unilateral renal pedicle clamping for 22 minutes followed by 6 days of recovery and contralateral nephrectomy, with sacrifice on day 7 (n=4), with sham-nephrectomy controls (n=3). The UUO model involved left ureter ligation for 4 days (n=4), with non-obstructed contralateral kidneys serving as controls (n=4). These models represent distinct injury mechanisms, specifically ischemic/reperfusion damage in AKI and obstructive nephropathy in UUO-CKD, enabling identification of both shared and disease-specific proteomic signatures.

**Figure 1.**
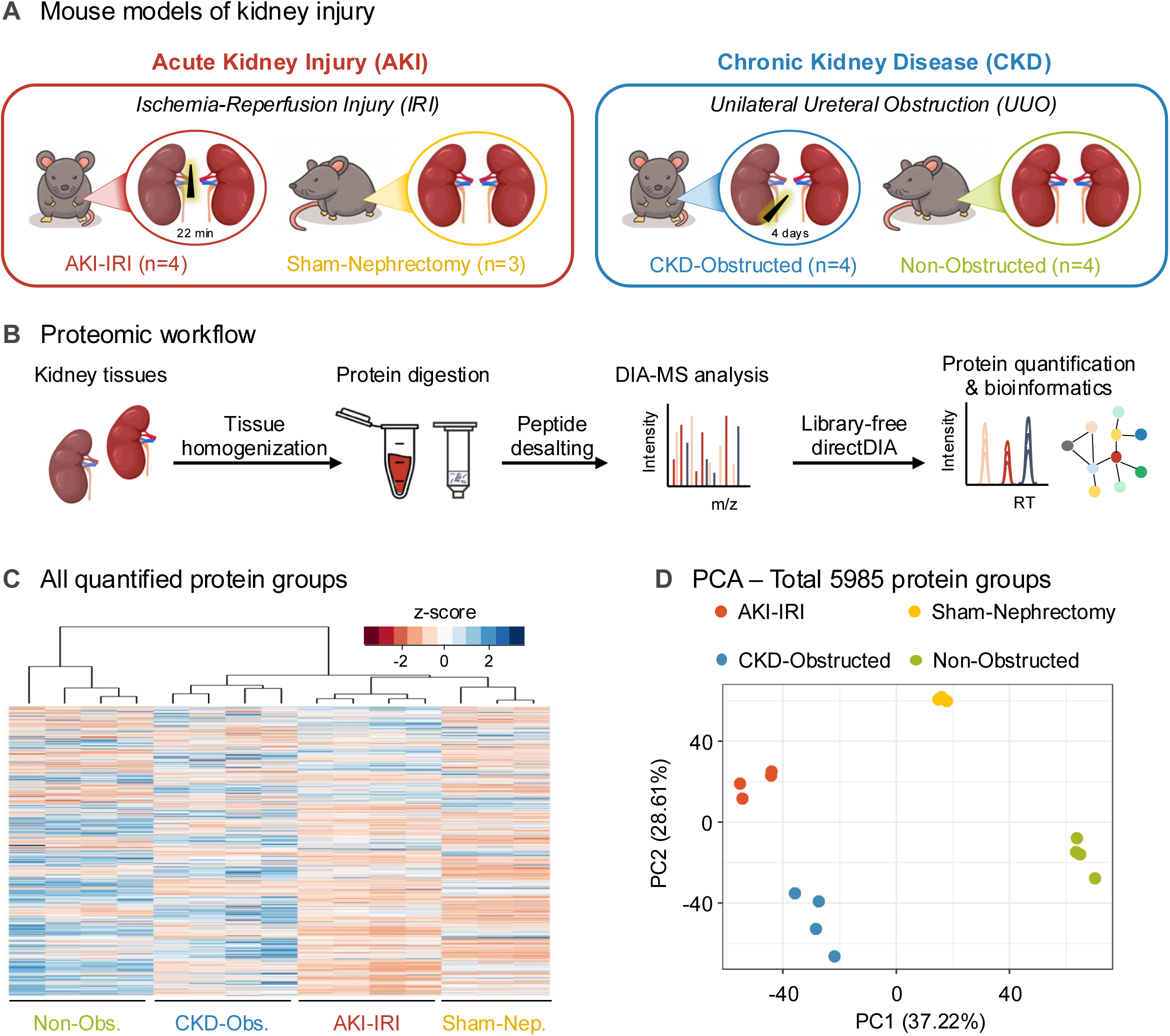
Study design and proteomic workflow. (A) Mouse models of kidney injury. Acute kidney injury (AKI) was induced by unilateral ischemia-reperfusion injury (IRI; 22-minute renal pedicle clamping, 7-day recovery; n=4) with sham-nephrectomy controls (n=3). Chronic kidney disease (CKD) was induced by unilateral ureteral obstruction (UUO; 4-day obstruction; n=4) with non-obstructed contralateral kidneys as controls (n=4). (B) Proteomic workflow: after kidney tissue homogenization, proteins were extracted, digested with trypsin, and desalted prior to data-independent acquisition mass spectrometry (DIA-MS) on an Orbitrap Eclipse Tribrid mass spectrometer, followed by library-free directDIA protein quantification and bioinformatic analysis. (C) Heatmap of scaled protein abundance (z-score) with hierarchical clustering of all 5,985 protein groups quantified with ≥2 unique peptides. (D) Principal component analysis (PCA) of all quantified protein groups demonstrated clear separation of all four experimental groups (PC1: 37.22%; PC2: 28.61%).

Kidney tissues were prepared for deep global proteomic analysis by tissue homogenization, protein extraction and digestion with trypsin **(Figure 1B)**. Samples were acquired by DIA-MS on an Orbitrap Eclipse Tribrid mass spectrometer, with directDIA data processing, which yielded quantification of 5,985 protein groups with high confidence (≥2 unique peptides; **Supplemental Table 1)**. Hierarchical clustering of all quantified protein groups demonstrated clear separation of all four experimental groups **(Figure 1C)**, and principal component analysis (PCA) confirmed this separation, with PC1 capturing 37.22% and PC2 capturing 28.61% of total variance **(Figure 1D)**, revealing robust experimental design and distinct molecular profiles between conditions.

### Differential expression analysis reveals extensive proteome remodeling

To identify protein changes associated with AKI and CKD, we performed differential analysis comparing each disease condition to its respective control. Differential expression analysis used stringent statistical criteria (q < 0.01, |Log2FC| > 0.58, corresponding to ≥1.5-fold change) and identified substantial proteome alterations in both disease models **(Figure 2A; Supplemental Table 1)**. AKI induction resulted in 2,729 significantly altered proteins (1,266 down-regulated, 1,464 up-regulated), representing 46% of the quantified proteome. The top down-regulated proteins were 4-hydroxyphenylpyruvate dioxygenase (Hpd), skeletal muscle isoform of Myosin light chain 1/3 (Myl1), Cytochrome P450 (Cyp2b10), S-adenosylhomocysteine hydrolase-like 2 (fragment, Ahcyl2), and Cytochrome P450 7B1 (Cyp7b1), while the top up-regulated proteins were WD repeat and FYVE domain containing 4 (Wdfy4), Collagen alpha-1(VIII) chain (Col8a1), C5a anaphylatoxin chemotactic receptor 1 (C5ar1), Toll-like receptor 2 (Tlr2), and Proteinase-activated receptor 1 (F2r) **(Figure 2B)**. Over-representation analysis (ORA) using ConsensusPathDB (48, 49) showed that proteins down-regulated in AKI were enriched for branched-chain amino acid metabolism, respiratory electron transport chain, cellular respiration, fatty acid oxidation, and peroxisome organization, whereas up-regulated proteins were enriched for antigen processing and presentation, ECM organization, nuclear export of ribonucleoprotein complexes, and immune processes **(Figure 2C; Supplemental Table 2)**. CKD showed 1,940 significantly altered proteins (1,068 down-regulated, 872 up-regulated), representing 32% of the quantified proteome **(Figure 2D)**. The proteins Solute carrier family 22 member 7 (Slc22a7), Phosphatidylinositol N-acetylglucosaminyltransferase subunit Q (Pigq), RUS family member 1 (Rusf1), Sodium-coupled monocarboxylate transporter 2 (Slc5a12), and Sodium-dependent neutral amino acid transporter B(0)AT1 (Slc6a19) showed the strongest down-regulation in CKD. In contrast, the most increased proteins were BRCA1-associated ATM activator 1 (Brat1), Cornifin-A (Sprr1a), Metallothionein-2 (Mt2), Uroplakin-2 (Upk2), and Major urinary protein 7 (fragment, Mup7) **(Figure 2E)**. ORA of the CKD signature revealed down-regulation of mitochondrial and metabolic processes similar to AKI, and enrichment of antigen processing and presentation, ECM organization, complement activation, coagulation, and immune responses among up-regulated proteins **(Figure 2F; Supplemental Table 2)**.

**Figure 2.**
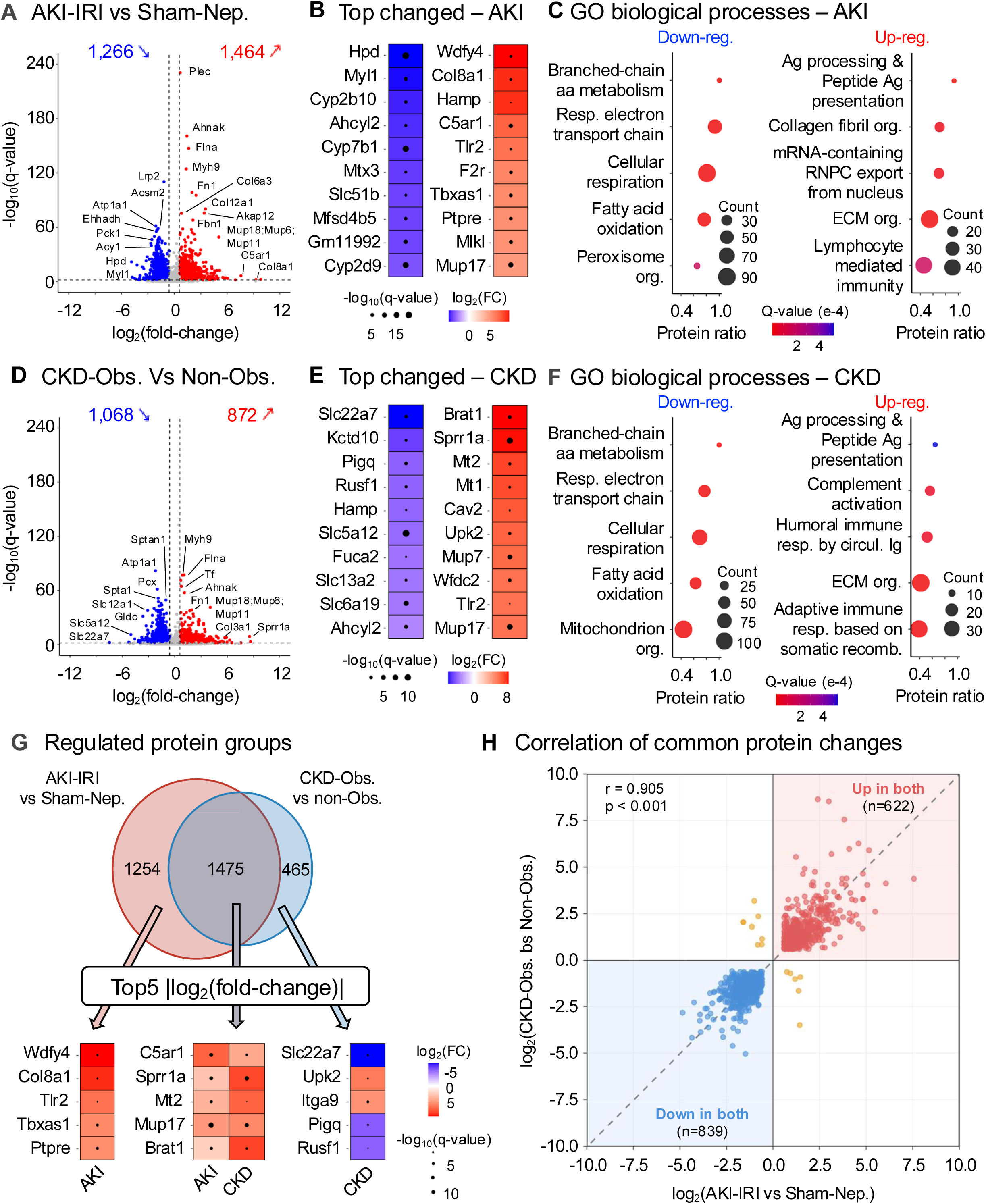
Differential analysis and proteome overlap in AKI and CKD. (A, D) Volcano plots of the protein groups quantified in (A) AKI (AKI-IRI vs Sham-Nephrectomy; 2,729 significantly altered: 1,266 down-regulated and 1,464 up-regulated) and (D) CKD (CKD-Obstructed vs Non-Obstructed; 1,940 significantly altered: 1,068 down-regulated and 872 up-regulated). Significance was defined as q-value < 0.01 and |log2(fold-change)| > 0.58. (B, E) Heatmaps of the log2(fold-change) of the top 10 down- and up-regulated proteins in (B) AKI and (E) CKD; dot size represents −log10(q-value). (C, F) Selected Gene Ontology (GO) biological processes from over-representation analysis of the down- and up-regulated proteins in (C) AKI and (F) CKD (ConsensusPathDB; q-value < 0.001, term_level = 4); full lists are provided in Supplemental Table 2. (G) Venn diagram of the AKI and CKD signatures (1,475 shared, 1,254 AKI-specific, and 465 CKD-specific protein groups), with heatmaps of the top 5 proteins with the largest |log2(fold-change)| in each subset. (H) Pearson correlation of the AKI and CKD log2(fold-change) for the 1,475 shared proteins (r = 0.905, p < 0.001). Red, common up-regulated (n=622); blue, common down-regulated (n=839); orange, opposite change direction (n=14). The dotted line represents x = y.

The greater number of altered proteins in AKI may reflect the acute, global nature of ischemic injury affecting the entire kidney, whereas the 4-day UUO model captures a more anatomically localized chronic fibrotic response. A striking finding emerged from overlap analysis: 1,475 proteins (46% of all altered proteins) were significantly changed in both AKI and CKD **(Figure 2G; Supplemental Table 3)**. This substantial overlap indicates a conserved core kidney injury response activated regardless of the initial insult mechanism. The shared proteome included 622 proteins up-regulated in both conditions and 839 proteins down-regulated in both conditions, and the log2(fold-change) values of the shared proteins were strongly correlated between AKI and CKD (Pearson r = 0.905, p < 0.001; **Figure 2H)**. Disease-specific changes included 1,254 proteins unique to AKI and 465 proteins unique to CKD **(Figure 2G)**.

### Shared and disease-specific biological processes

ORA of the shared proteins revealed the biological significance of the conserved core injury response **(Figure 3C; Supplemental Table 4; Supplemental Figure 1)**. The 622 shared up-regulated proteins were enriched for immune system activation, extracellular matrix (ECM) organization, inflammatory response, DNA-binding transcription factor activity, circulatory system development, and cell migration, indicating that both acute ischemic injury and chronic obstructive injury trigger common inflammatory and tissue remodeling programs.

**Figure 3.**
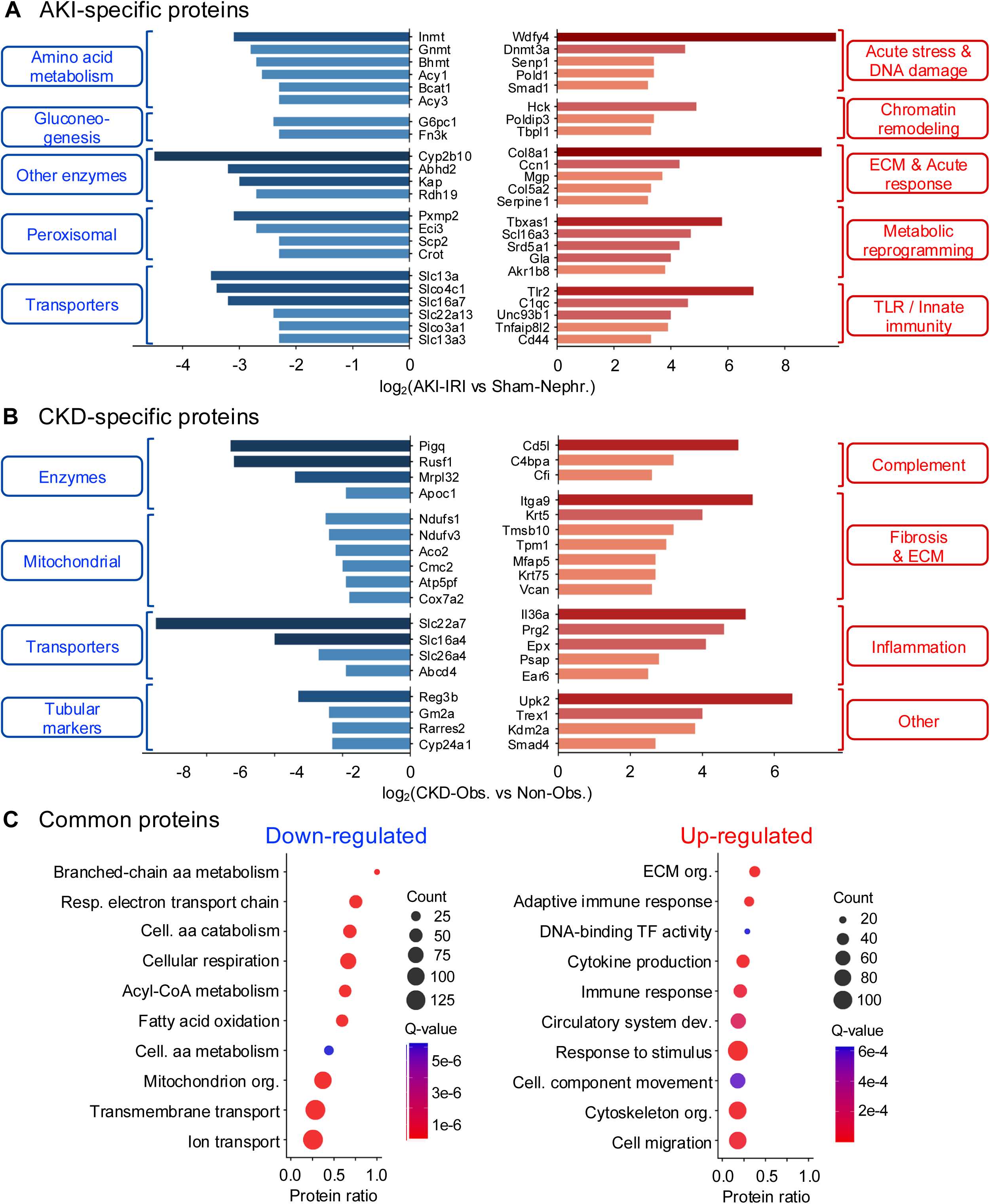
Disease-specific and shared proteome signatures in AKI and CKD. (A, B) Bar charts of selected top down-regulated (blue, left) and up-regulated (red, right) proteins specifically altered in (A) AKI (AKI-IRI vs Sham-Nephrectomy; 1,254 protein groups: 418 down-regulated and 836 up-regulated) or (B) CKD (CKD-Obstructed vs Non-Obstructed; 465 protein groups: 223 down-regulated and 242 up-regulated), grouped by biological function. Values shown are log2(fold-change). (C) Selected GO biological processes from over-representation analysis of the 839 common down-regulated (left) and 622 common up-regulated (right) proteins (ConsensusPathDB; q-value < 0.001, term_level = 4); full lists are provided in **Supplemental Table 4**.

Remarkably, the 839 shared down-regulated proteins were dominated by mitochondrial processes **(Figure 3C)**, consistent with the near-identical down-regulated processes observed for AKI and CKD individually **(Figure 2C and 2F)**. Key down-regulated processes included cellular respiration, oxidative phosphorylation, fatty acid oxidation, branched-chain amino acid metabolism, and transmembrane and ion transport. This conserved metabolic collapse signature, affecting the same pathways regardless of whether injury is acute or chronic, represents a fundamental and striking feature of kidney injury pathophysiology. The kidney’s exceptionally high metabolic demand makes it uniquely vulnerable to mitochondrial dysfunction, and our data demonstrate that this vulnerability is exploited by diverse injury mechanisms.

Disease-specific signatures revealed distinct pathophysiological features **(Figure 3A-B; Supplemental Table 3)**. The AKI-specific up-regulated proteins (n=836) were involved in chromatin organization (e.g., Polymerase delta-interacting protein 3, Poldip3), DNA damage response (e.g., DNA (cytosine-5)-methyltransferase 3A, Dnmt3a), ECM organization (e.g., Collagens Col8a1 and Col5a2; CCN family member 1, Ccn1), innate immunity (e.g., Complement C1q subcomponent subunit C, C1qc), and metabolic reprogramming (e.g., Alpha-galactosidase A, Gla), reflecting acute cellular stress responses to ischemia-reperfusion. In contrast, proteins related to amino acid metabolism (e.g., Indolethylamine N-methyltransferase, Inmt), gluconeogenesis (e.g., Glucose-6-phosphatase catalytic subunit 1, G6pc1), members of the solute carrier (Slc) families, and peroxisomal proteins (e.g., Enoyl-CoA delta isomerase 3, Eci3; Carnitine O-octanoyltransferase, Crot) were specifically down-regulated in AKI (n=418).

With respect to CKD, the specifically down-regulated proteins (n=223) included numerous transporters from the Slc families, mitochondrial proteins (e.g., Aconitase 2, Aco2), and renal chemerin (Rarres2), known to be associated with CKD (50). The CKD-specific up-regulated proteins (n=242) included sustained fibrosis markers (e.g., Tropomyosin alpha-1 chain, Tpm1, previously reported to play a role in the initiation of kidney fibrosis (51)), complement cascade components (e.g., Complement factor I, Cfi), and inflammatory proteins (e.g., Eosinophil peroxidase, Epx), signatures of chronic tissue remodeling and progressive scarring that define the chronic injury phenotype.

### Identification of AKI-to-CKD progression markers

A major goal of this study was to identify proteins that serve as markers of disease progression from acute to chronic kidney injury. We identified 95 candidate progression markers, proteins significantly up-regulated in both AKI and CKD but with a fold-change at least 1.5-fold higher in the chronic condition **(Figure 4A; Supplemental Figure 2; Supplemental Table 3)**. These proteins span six functional categories: stress response and metal homeostasis (Sprr1a, Mt2, Sfn, Grn), DNA damage and transcription (Brat1, Polr2i, Snrpc), ECM and fibrosis (Col3a1, Actn1, Fndc3b, Krt14), signaling and phosphorylation (Ppp2r5c, Fyb1, Riok3, Shc1), coagulation and complement (Ifi30, F10, F9), and lipid metabolism (Agpat4, Rxrb, Xdh).

**Figure 4.**
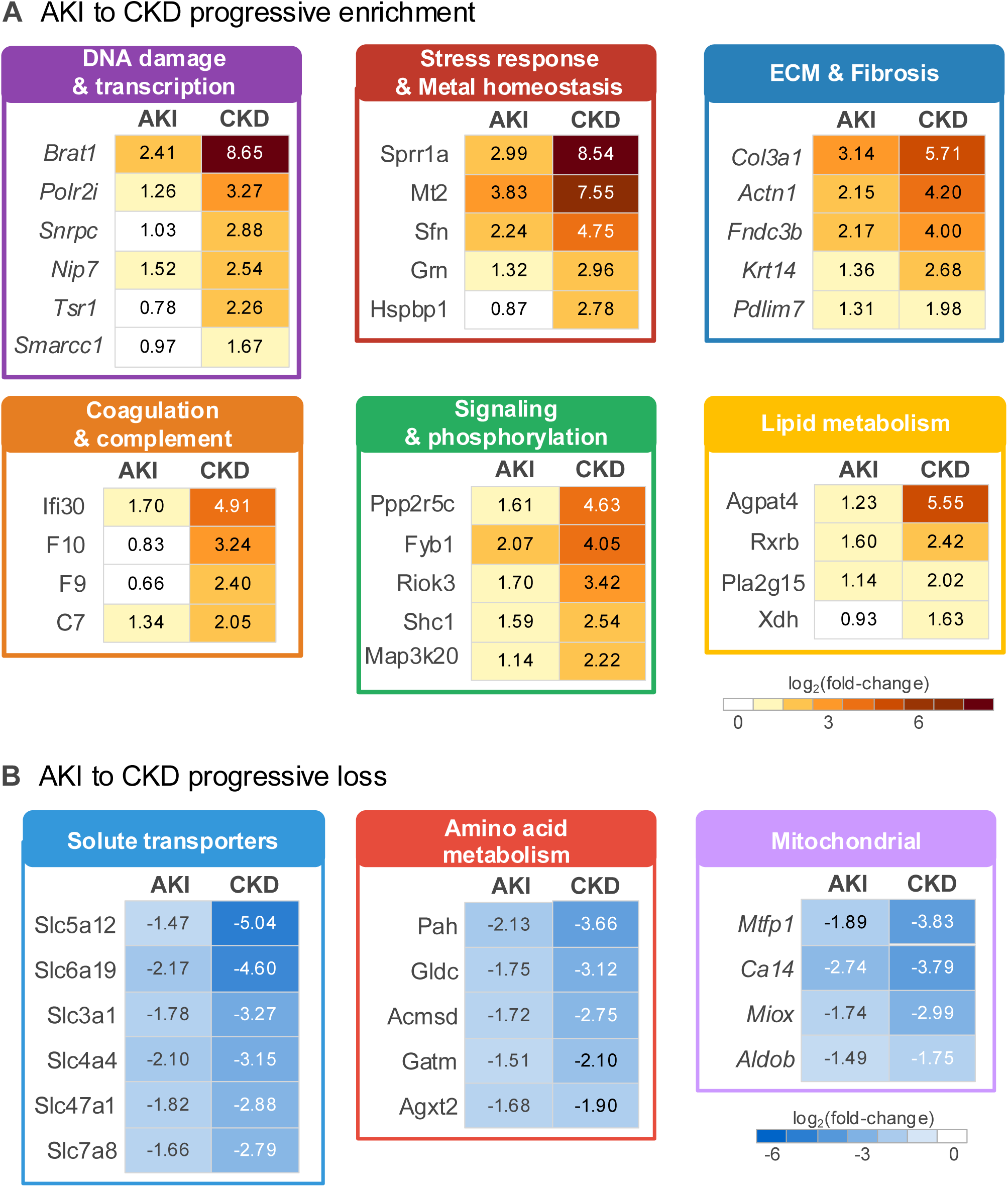
Identification of candidate AKI-to-CKD transition markers. (A) Candidate progression markers with progressive gain from AKI to CKD: 95 protein groups were significantly up-regulated in both AKI and CKD, with a fold-change at least 1.5-fold higher in CKD (CKD-Obstructed vs Non-Obstructed) than in AKI (AKI-IRI vs Sham-Nephrectomy). (B) Proteins with progressive loss from AKI to CKD: 93 protein groups were significantly down-regulated in both AKI and CKD, with a fold-change at least 1.5-fold lower in CKD than in AKI. Heatmaps show the log2(fold-change) of selected proteins, grouped by biological function or cellular compartment. Full lists are provided in **Supplemental Table 3**.

The progression marker Brat1 showed the most dramatic escalation, increasing from Log2FC +2.41 in AKI to Log2FC +8.65 in CKD, a 76-fold increase in expression from acute to chronic kidney injury. Brat1 (BRCA1-associated ATM activator 1) functions in DNA damage response and chromatin remodeling, suggesting that persistent genotoxic stress may be a driver of CKD progression. Similarly, the fibrosis marker Col3a1 (Collagen type III alpha 1) increased from Log2FC +3.14 in AKI to +5.71 in CKD, consistent with progressive ECM deposition during chronic injury. Because the AKI and CKD signatures were derived from two distinct injury models rather than a single longitudinal time course, we consider these proteins candidate progression markers. These candidate progression markers represent potential biomarkers for identifying patients at risk of AKI-to-CKD transition and highlight targetable pathways for therapeutic intervention.

Complementary to the escalating progression markers, we identified 93 proteins showing progressive loss with down-regulation in both AKI and CKD but significantly more suppressed in CKD (fold-change at least 1.5-fold lower in CKD than in AKI; **Figure 4B)**. These progressive loss proteins included solute transporters (Slc5a12 at −1.5/−5.0, Slc6a19 at −2.2/−4.6), ion channels (Kcnj16 at −2.7/−4.1), amino acid metabolism enzymes, and multiple mitochondrial proteins (e.g., Mtfp1, Ca14, and Miox).

### Opposite directionality in proteins for AKI and CKD reveals phase-specific biology

We identified 14 proteins with divergent expression patterns, meaning that they were up-regulated in one condition but down-regulated in the other **(Figure 2H and Figure 5; Supplemental Table 3)**. Six proteins were up-regulated in AKI and down-regulated in CKD, and eight proteins showed the reverse pattern. These “opposite direction” proteins provide unique insights into the distinct biology of acute versus chronic kidney injury and represent potential diagnostic markers to distinguish disease phases. *Mfn2* (Mitofusin-2) showed up-regulation in AKI (+1.37 Log2FC) but down-regulation in CKD (−1.65 Log2FC). Mfn2 is essential for mitochondrial fusion, and its acute up-regulation likely represents a compensatory response to maintain mitochondrial network integrity following ischemic injury. The subsequent down-regulation in CKD suggests a shift toward mitochondrial fragmentation and dysfunction during chronic disease, consistent with the metabolic collapse signature observed in our pathway analysis.

**Figure 5.**
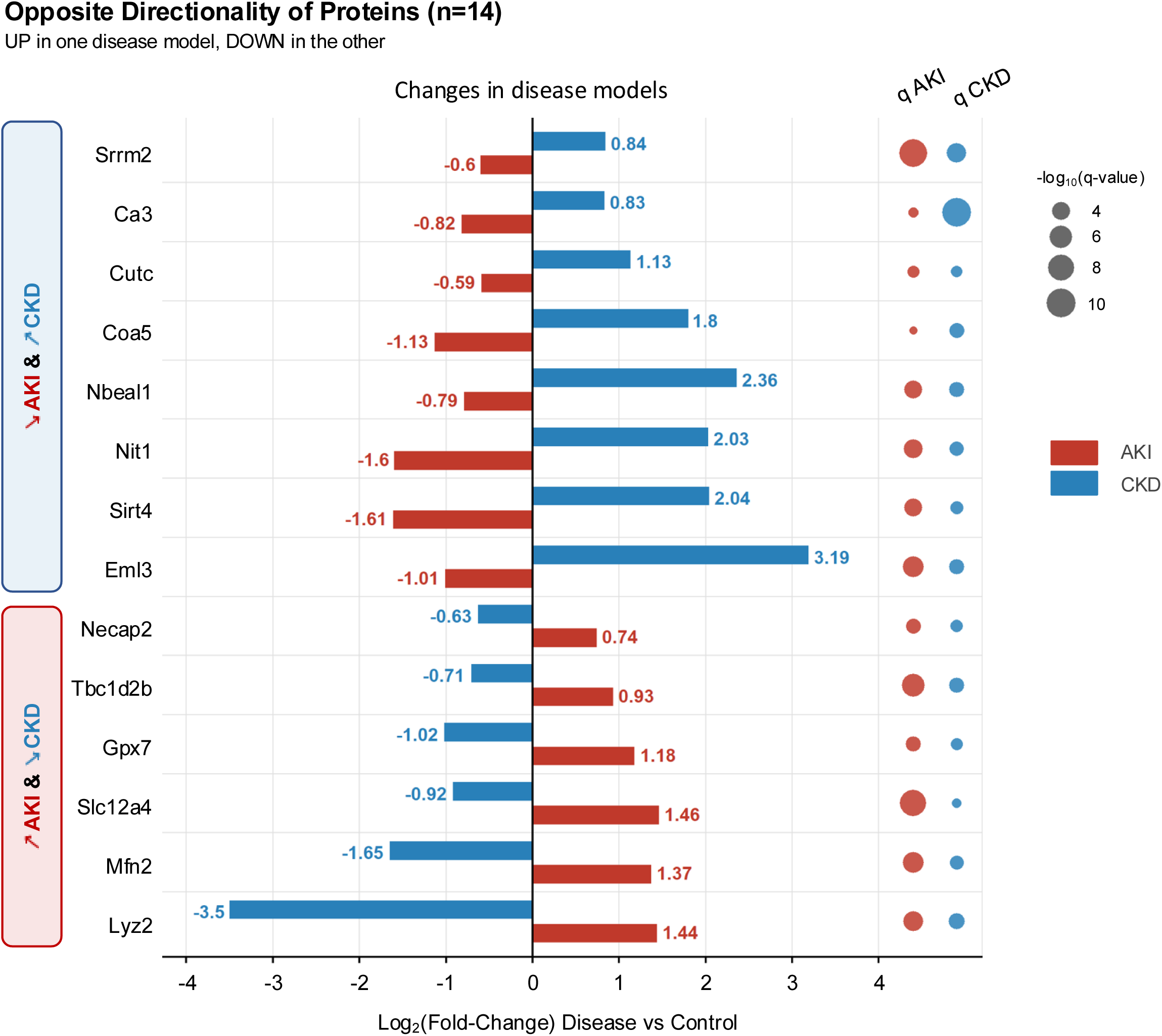
Proteins with opposite change direction in AKI and CKD. Of the 1,475 protein groups common to the AKI and CKD signatures, 14 showed opposite change directions: six were up-regulated in AKI and down-regulated in CKD, and eight showed the reverse pattern. The bar chart represents the log2(fold-change) in AKI (AKI-IRI vs Sham-Nephrectomy, red) and CKD (CKD-Obstructed vs Non-Obstructed, blue), and dot size represents −log10(q-value).

*Lyz2* (Lysozyme C-2) exhibited the most dramatic opposite pattern: up-regulated in AKI (+1.44) but strongly down-regulated in CKD (−3.50), indicating acute inflammatory response followed by immune exhaustion during chronic disease. Conversely, *Sirt4* (Sirtuin-4) was down-regulated in AKI (−1.61) but up-regulated in CKD (+2.04), suggesting metabolic stress acutely followed by adaptive metabolic reprogramming chronically. Similarly, *Eml3* (EMAP-like 3), a microtubule-associated protein, was down-regulated in AKI (−1.01) but strongly up-regulated in CKD (+3.19), indicating cytoskeletal disruption acutely followed by reorganization associated with fibrosis. These phase-specific markers represent compelling diagnostic tools to distinguish acute from chronic kidney injury in clinical settings where this distinction can be challenging.

### Co-expression network analysis identifies disease-associated modules

To identify coordinated protein networks associated with each disease state, we performed weighted gene co-expression network analysis (WGCNA) (52) on the abundance of all quantified protein groups **(Figure 6; Supplemental Tables 5 and 6)**. This identified 27 modules ranging from 817 proteins (Turquoise) to 61 proteins (Dark orange) **(Figure 6A-B)**. Three modules showed significant positive correlation with AKI (r > 0.8, p < 0.05): Dark-red (103 proteins), Light-cyan (125 proteins), and Royal-blue (103 proteins) **(Supplemental Figure 3A)**, and Kruskal-Wallis testing of the module eigengenes confirmed their increase in AKI compared to CKD and controls **(Supplemental Figure 3B)**. Module Dark-red was enriched for chromatin and chromosome organization, consistent with the acute stress response to ischemia-reperfusion. Module Light-cyan showed enrichment for ECM organization, and Module Royal-blue for inflammatory response pathways **(Supplemental Table 6)**.

**Figure 6.**
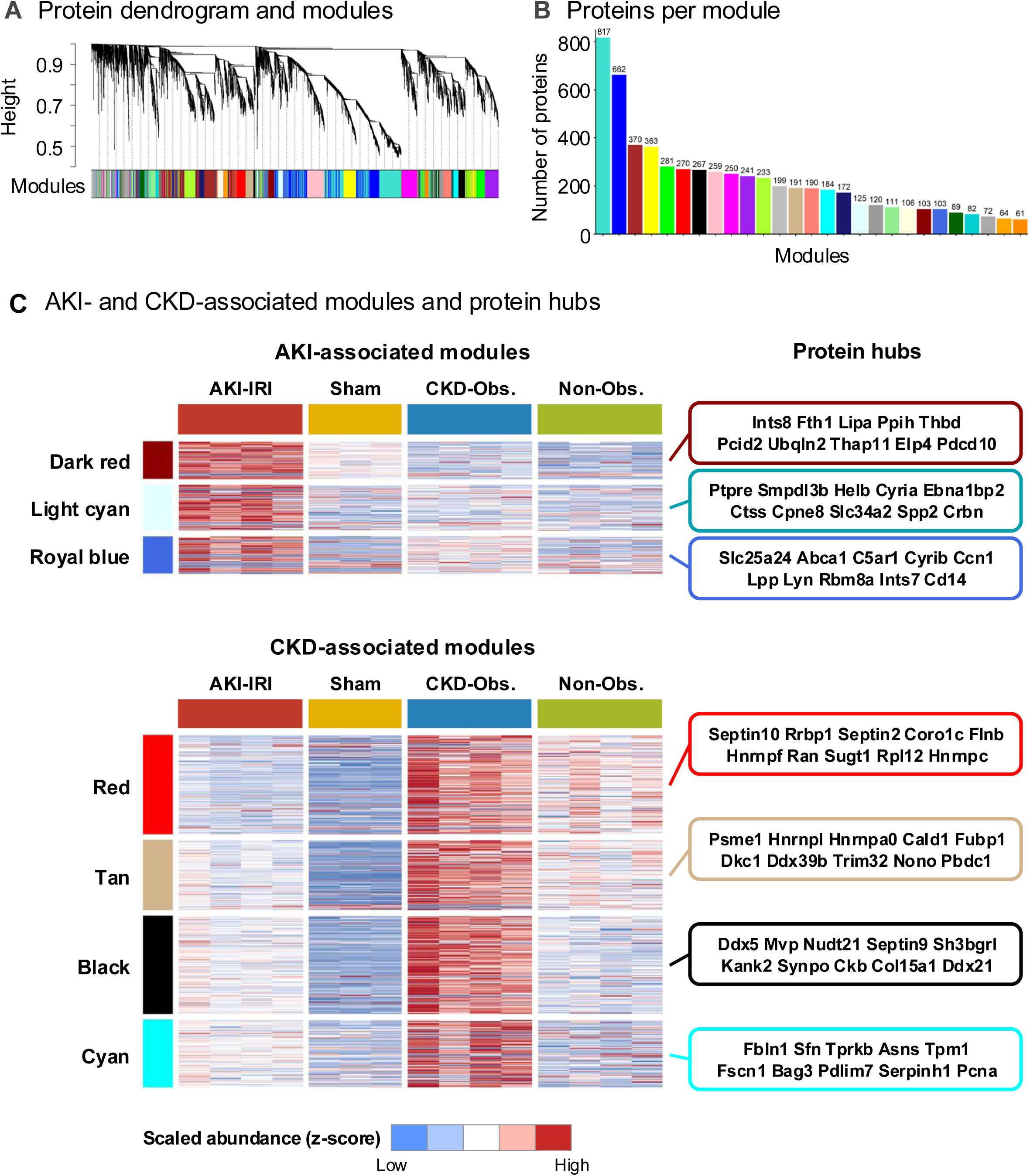
Identification of AKI- and CKD-associated modules by protein co-expression network analysis. Weighted gene co-expression network analysis (WGCNA) was conducted on all 5,985 quantified protein groups (52). (A) Protein cluster dendrogram and identified modules. (B) Number of proteins in each of the 27 identified modules. (C) Heatmaps of scaled protein abundance across replicates of the four experimental groups for the three AKI-associated modules (Dark red, Light cyan, and Royal blue; top) and the four CKD-associated modules (Red, Tan, Black, and Cyan; bottom) (r > 0.8, p < 0.05; Supplemental Figure 3), with the protein hubs showing the strongest protein-module kME correlations.

Four modules were significantly associated with CKD: Red (270 proteins), Tan (191 proteins), Black (267 proteins), and Cyan (184 proteins) **(Supplemental Figure 3A and 3B)**. Module Red was enriched for transcription and protein synthesis, Module Tan for ribonucleoprotein complexes, Module Black for nuclear proteins, and Module Cyan for the response to stress. The top protein hubs, based on protein-module kME correlations, were Integrator complex subunit 8 (Ints8) for Dark-red, Receptor-type tyrosine-protein phosphatase epsilon (Ptpre) for Light-cyan, and Mitochondrial adenyl nucleotide antiporter Slc25a24 for Royal-blue; and Septin-10 for Red, Proteasome activator complex subunit 1 (Psme1) for Tan, Probable ATP-dependent RNA helicase Ddx5 for Black, and Fibulin-1 (Fbln1) for Cyan **(Figure 6C; Supplemental Table 5)**. The identification of these disease-specific protein networks provides a systems-level framework for understanding the coordinated molecular responses underlying acute versus chronic kidney injury.

### Subcellular localization analysis reveals broad metabolic enzyme loss in AKI and CKD

To highlight the subcellular distribution of differentially abundant proteins, we mapped our findings onto a LOPIT (Localization of Organelle Proteins by Isotope Tagging) reference dataset **(Figure 7; Supplemental Figure 4)** (53, 54). This analysis revealed prominent up-regulation of proteins localized to the nucleus (both chromatin- and non-chromatin-associated proteins) and, to a lesser extent, to the lysosome and the endoplasmic reticulum/Golgi apparatus in both conditions, with extensive and coordinated down-regulation of mitochondrial and peroxisomal proteins. Notably, 353 down-regulated proteins in CKD mapped to mitochondria, compared to only 4 up-regulated mitochondrial proteins, a striking asymmetry underscoring the severity of mitochondrial dysfunction in chronic kidney disease.

**Figure 7.**
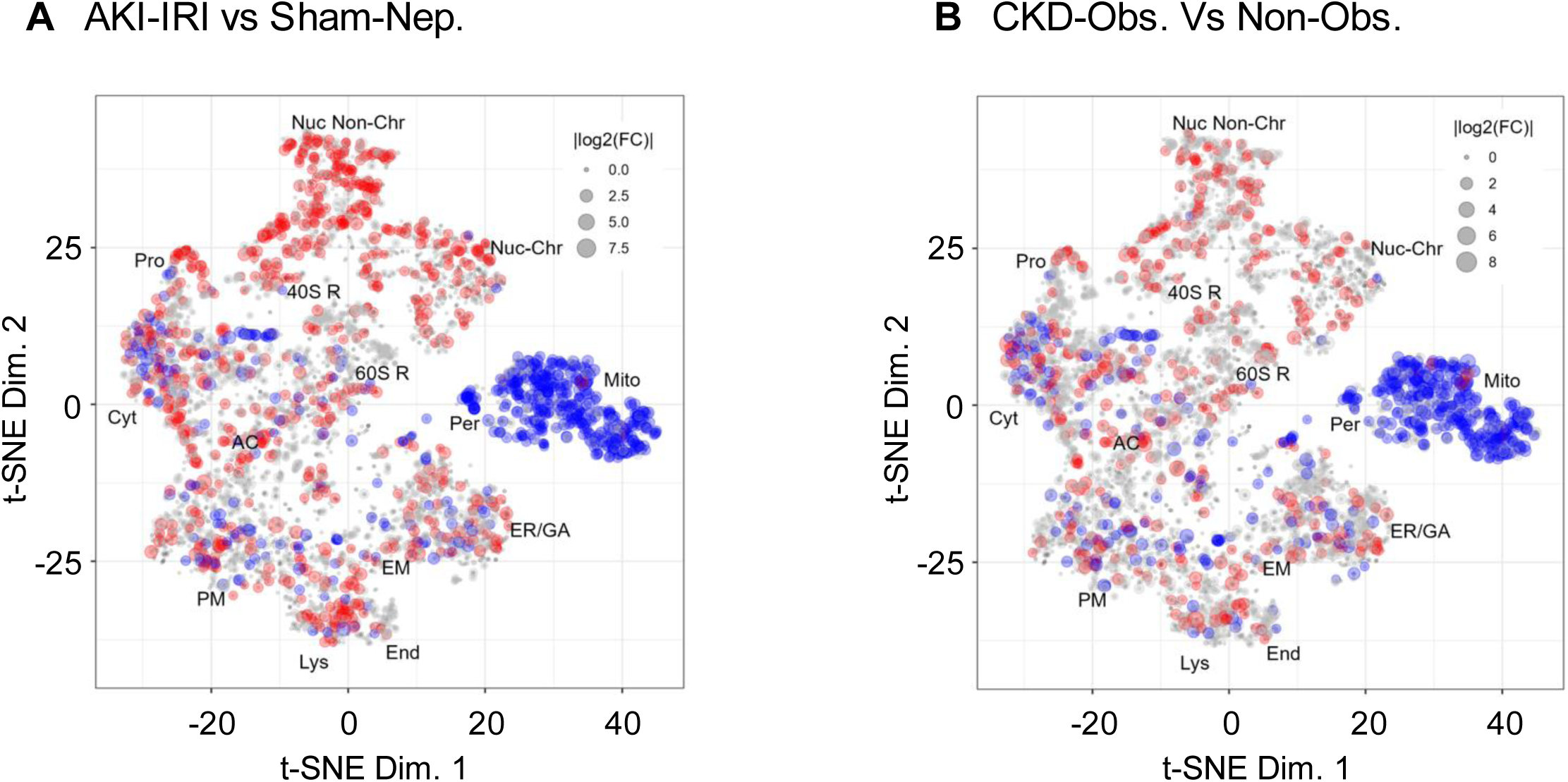
Coordinated loss of mitochondrial and peroxisomal proteins in AKI and CKD. LOPIT map representing the subcellular distribution of the proteins altered in (A) AKI and (B) CKD. Dots represent the log2(fold-change) of each protein, with up-regulation displayed in red and down-regulation in blue. Assigned coordinates are shown for 2,818 proteins, including 1,251 altered in AKI and 890 altered in CKD, for the 40S ribosome (40S R), 60S ribosome (60S R), actin cytoskeleton (AC), cytosol (Cyt), endoplasmic reticulum/Golgi apparatus (ER/GA), endosome (End), extracellular matrix (EM), lysosome (Lys), mitochondria (Mito), nucleus chromatin (Nuc-Chr), nucleus non-chromatin (Nuc Non-Chr), peroxisome (Per), plasma membrane (PM), proteasome (Pro), and undefined (grey) subcellular locations, based on the localization of proteins determined in (54). All significantly altered proteins mapped to the peroxisome were down-regulated in both AKI (n=18) and CKD (n=12). Down-regulated peroxisomal proteins include Acox1, Acox2, Acox3, Ehhadh, Hsd17b4, Acaa1a, Scp2, Crot, Crat, Amacr, Pecr, Cat, Pex1, Pex13, Pex14, and Pxmp2 (Supplemental Table 7).

A particularly noteworthy finding from the LOPIT analysis was the widespread down-regulation of proteins localized to the peroxisome in both AKI and CKD **(Figure 7)**. In the LOPIT map, 18 significantly altered peroxisomal proteins in AKI and 12 in CKD were all down-regulated, and no up-regulated protein mapped to the peroxisome. To capture the peroxisomal proteome beyond the LOPIT coverage, we examined all quantified protein groups with established peroxisomal localization **(Supplemental Table 7)**. Of these, 45 protein groups were significantly down-regulated in AKI and 28 in CKD, and none were up-regulated in either model. The down-regulated peroxisomal proteins spanned the β-oxidation machinery (Acox1, Acox2, Acox3, Ehhadh, Hsd17b4, Acaa1a, Acaa1b, Scp2, Amacr, Pecr, Decr2, Ech1, and Eci3), carnitine and acyl-CoA metabolism (Crot, Crat, Acot4, Acot8, Nudt7, Nudt12, Nudt19, Mlycd, and Acnat1), α-oxidation (Phyh and Hacl1), oxidases and antioxidant defense (Cat, Dao, Hao2, and Pipox), ether lipid synthesis (Agps, Gnpat, and Far1), and peroxisome biogenesis, import, and membrane transport (Pex1, Pex6, Pex7, Pex11a, Pex13, Pex14, Pex16, Abcd3, Pxmp2, Pxmp4, and Mpv17l). Twenty-eight peroxisomal protein groups were down-regulated in both models, and for 27 of them the decrease was larger in AKI than in CKD; for example, the rate-limiting enzyme of peroxisomal β-oxidation Acox1 decreased by log2(FC) = −2.91 in AKI and −1.63 in CKD, and the L-bifunctional enzyme Ehhadh by −1.96 and −0.77, respectively. Peroxisomal depletion is therefore a shared and early feature of acute and chronic kidney injury rather than a CKD-specific process, consistent with an independent IRI proteomics study reporting defective peroxisomal fatty acid oxidation during the progression of AKI and repair (55). Together, these findings reveal a broad collapse of proximal tubule metabolic capacity across mitochondria and peroxisomes in both AKI and CKD, linking peroxisomal dysfunction to tubular lipotoxicity and fibrotic progression (56, 57).

### Senescence-associated secretory signature in AKI and CKD

Given the central role of cellular senescence in kidney aging and in the AKI-to-CKD transition, we interrogated our data with a curated panel of 116 senescence and SASP-associated proteins compiled from the SASP Atlas (58) and the SenMayo gene set (59) (Figure 8; Supplemental Table 8). Of the 54 panel proteins quantified, 40 proteins (41 protein groups) were significantly altered in at least one model, and nearly all of them were up-regulated (34 of 36 protein groups in AKI and 30 of 31 in CKD). Twenty-five proteins were up-regulated in both AKI and CKD (Figure 8A), including the tubular injury and SASP-associated factors Hepatitis A virus cellular receptor 1 (Havcr1, also known as KIM-1; log2(FC) = +4.83 in AKI and +3.26 in CKD), Osteopontin (Spp1; +4.24 and +3.26), Insulin-like growth factor-binding protein 7 (Igfbp7; +1.95 and +1.10), Urokinase-type plasminogen activator (Plau; +1.93 and +2.42), Vascular cell adhesion protein 1 (Vcam1; +2.59 and +3.29), Intercellular adhesion molecule 1 (Icam1), and Fibronectin (Fn1). Lysosomal proteins associated with the expanded lysosomal compartment of senescent cells were also consistently increased, including Cathepsins B, D, S, and Z (Ctsb, Ctsd, Ctss, and Ctsz), together with Galectin-1 and Galectin-3 (Lgals1 and Lgals3), S100a4, S100a6, Annexins A1 and A2 (Anxa1 and Anxa2), Clusterin (Clu), Vimentin (Vim), Stat1, and Gamma-interferon-inducible lysosomal thiol reductase (Ifi30; +1.70 and +4.91). Ten proteins were altered only in AKI (Figure 8B), including Plasminogen activator inhibitor 1 (Serpine1; +3.25), CCN family member 1 (Ccn1; +4.35), Cd44 (+3.25), and Neutrophil gelatinase-associated lipocalin (Lcn2; +1.07), whereas five proteins were altered only in CKD (Figure 8C), most notably High mobility group protein B2 (Hmgb2; +1.78). The canonical cell-cycle arrest markers p16 (Cdkn2a) and p21 (Cdkn1a), the DNA damage protein p53, and most low-abundance cytokines, including Il6 and Il1b, were not detected. Together, these data reveal a SASP-like secretory and lysosomal signature that is shared by acute and chronic kidney injury and is more pronounced in AKI.

**Figure 8.**
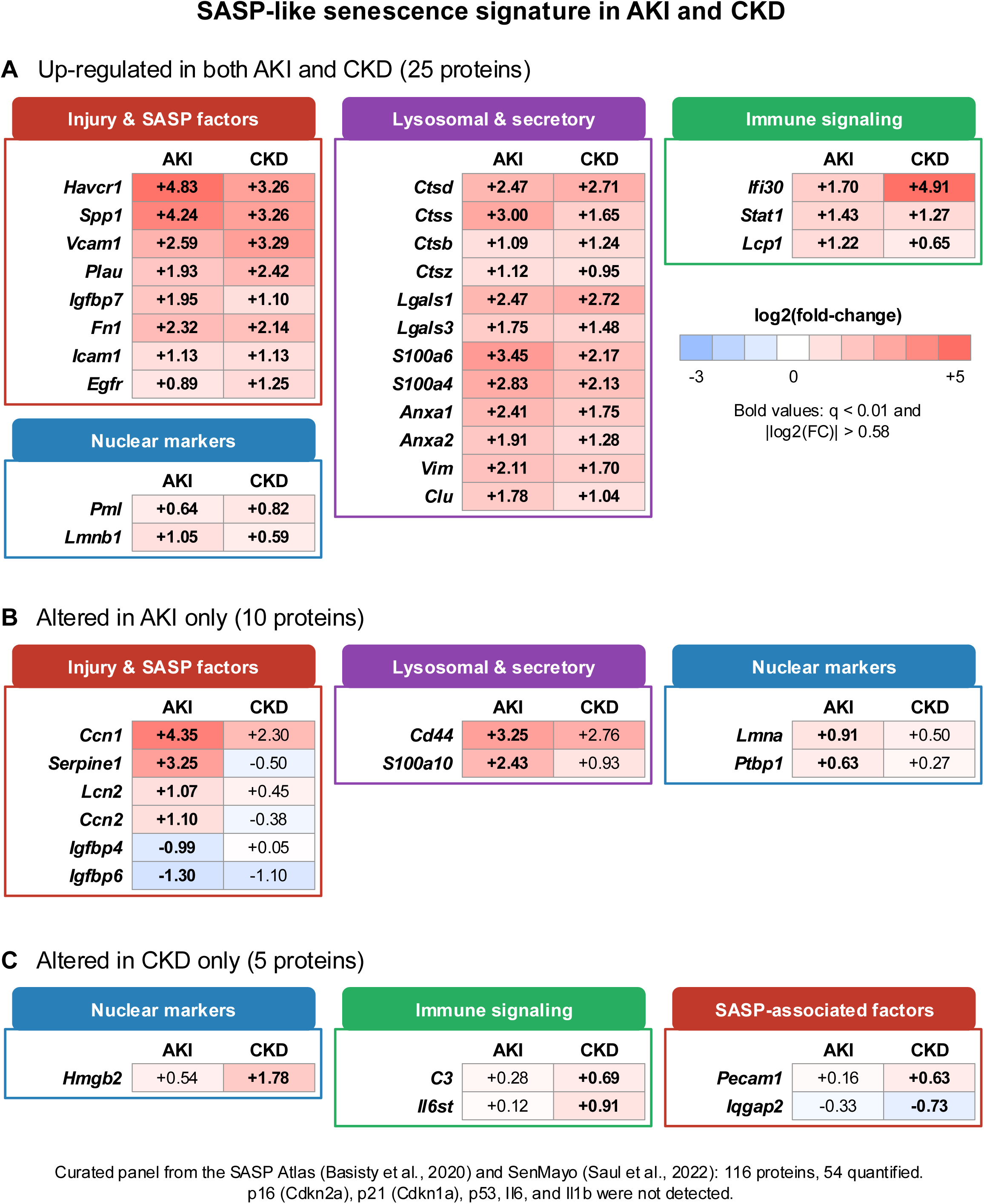
Senescence-associated secretory phenotype (SASP)-like signature in AKI and CKD. Heatmaps of the log2(fold-change) in AKI (AKI-IRI vs Sham-Nephrectomy) and CKD (CKD-Obstructed vs Non-Obstructed) for senescence and SASP-associated proteins from a curated panel based on the SASP Atlas (58) and the SenMayo gene set (59) that were significantly altered (q-value < 0.01, |log2(fold-change)| > 0.58) in (A) both AKI and CKD (25 proteins), (B) AKI only (10 proteins), or (C) CKD only (5 proteins), grouped by category. Bold values indicate significance. The canonical cell-cycle arrest markers p16 (Cdkn2a) and p21 (Cdkn1a) were not detected. Full panel results are provided in Supplemental Table 8.

### Integrative view of acute and chronic kidney injury proteomes

Our comprehensive proteomic analysis reveals both shared and distinct molecular signatures that advance understanding of kidney injury pathophysiology **(Figure 9)**. The finding that 46% of altered proteins are shared between AKI and CKD (despite different etiologies) indicates a conserved core kidney injury response. This core response is characterized by immune activation, ECM remodeling, and stress response programs on the up-regulated side, and by metabolic dysfunction on the down-regulated side, particularly mitochondrial impairment and the loss of solute transporters and peroxisomal proteins.

**Figure 9.**
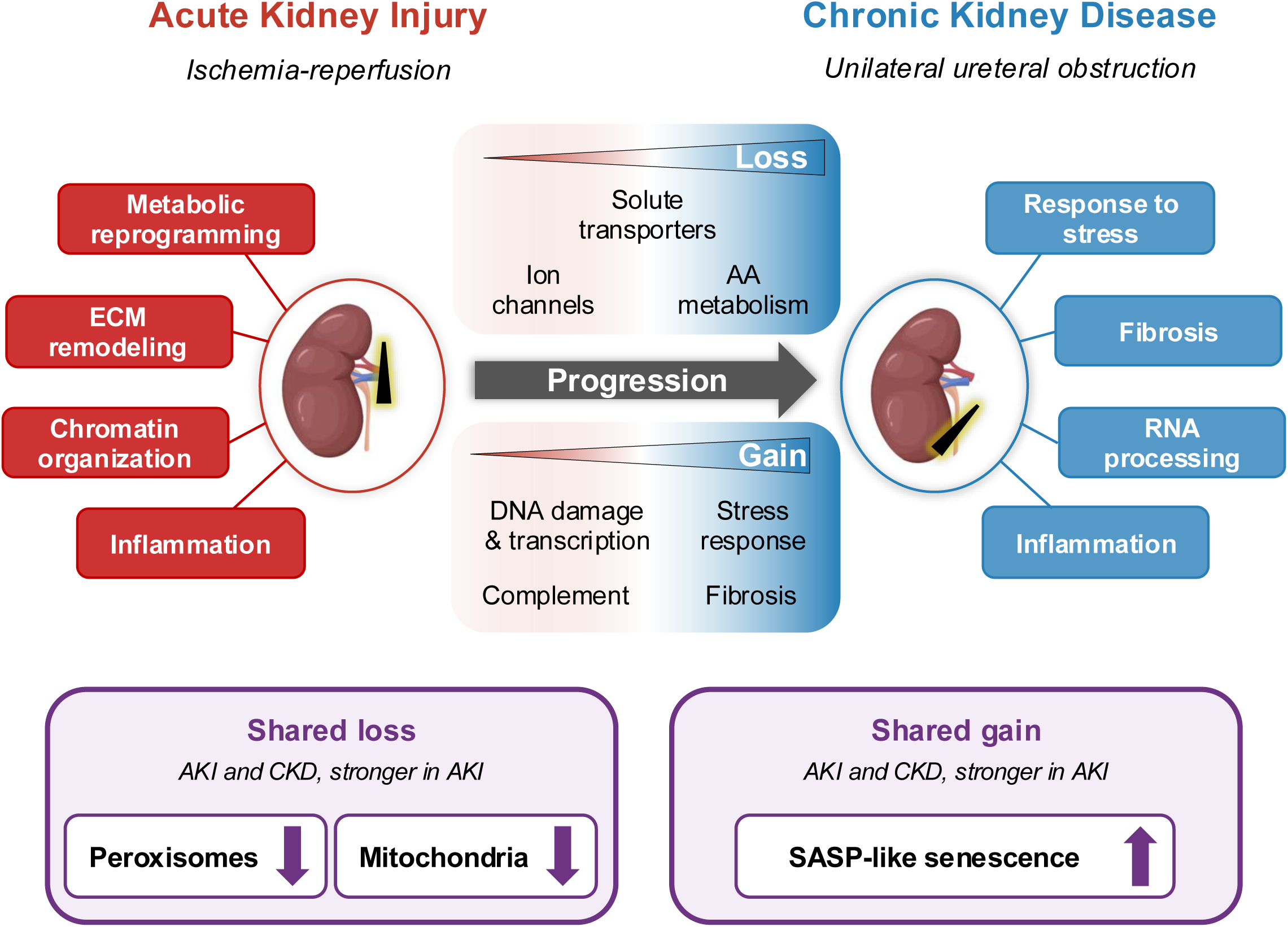
Concluding summary of the comparative proteomic analysis of AKI and CKD. Integrated summary schematic of the AKI and CKD proteomic signatures. Representative pathways specific to the AKI-IRI model are displayed in red and those specific to the CKD-UUO model in blue. Candidate progression markers with greater changes in CKD than in AKI are summarized in the center, with progressively depleted proteins (solute transporters, ion channels, and amino acid metabolism) above the arrow and progressively enriched proteins (DNA damage and transcription, stress response, complement, and fibrosis) below the arrow. The lower boxes show changes shared by both models and stronger in AKI: the loss of peroxisomal and mitochondrial proteins and a SASP-like senescence signature.

The near-identical metabolic collapse signature in both AKI and CKD is particularly noteworthy. The kidney has among the highest mitochondrial content of any organ, reflecting its enormous energy demands for active transport. Our data demonstrate that diverse injury mechanisms converge on mitochondrial and peroxisomal dysfunction as common pathogenic endpoints, with therapeutic implications: interventions targeting metabolic health may benefit patients with kidney injury regardless of the underlying cause.

The identification of candidate progression markers, progressive loss proteins, and opposite-direction proteins provides new tools for understanding and potentially predicting AKI-to-CKD transition. The 95 candidate progression markers may serve as biomarkers of disease progression, the 93 progressive loss proteins reveal the metabolic and functional consequences of chronicity, and the 14 opposite-direction proteins could help distinguish disease phases in clinical settings. In addition, the coordinated loss of peroxisomal proteins and the SASP-like signature observed in both models point to peroxisomal dysfunction and cellular senescence as shared processes of acute and chronic kidney injury. Importantly, several of these markers, including Mfn2, Sirt4, Brat1, and Acox1, represent potentially targetable pathways for therapeutic intervention aimed at preventing or slowing the transition from acute injury to chronic kidney disease.

## Limitations

Several limitations should be considered when interpreting these findings. First, while the IRI and UUO models are well-established, they represent only two of many potential kidney injury etiologies; the generalizability of our findings to other causes of AKI and CKD (e.g., nephrotoxic injury, diabetic nephropathy, glomerulonephritis) requires further investigation. Second, our analysis captures single time points (7 days post-IRI, 4 days post-UUO), and temporal dynamics of protein expression during injury and recovery warrant additional study. UUO represents an accelerated fibrosis model that captures key molecular features of CKD-associated remodeling, though it does not recapitulate the prolonged timeline of human CKD progression. Third, while whole-kidney proteomics provides comprehensive coverage, it cannot resolve cell-type-specific changes; single-cell or spatial proteomics approaches would complement these findings. Fourth, our study exclusively examined male mice, but female mice should also be investigated to determine the relevance of these findings. Fifth, canonical senescence markers such as p16 (Cdkn2a) and p21 (Cdkn1a), as well as most low-abundance cytokines, were not detected by our tissue DIA-MS workflow; the SASP-like signature therefore requires confirmation with orthogonal senescence assays. Finally, validation of candidate biomarkers and therapeutic targets in human samples and clinical cohorts will be essential for translation.

## Discussion

Our findings complement a growing body of single-cell transcriptomics literature characterizing cellular responses to AKI and the AKI-to-CKD transition. For example, published single-nucleus RNA-sequencing studies from the same IRI mouse model have primarily characterized inflammatory and profibrotic cell states, notably the failed-repair proximal tubule cell (FR-PTC) population defined by NF-κB activation (60–62). Interestingly, the reported FR-PTCs downregulate proximal tubule differentiation markers including *Slc5a12*, *Slc22a30*, and *Slc7a13* (61), which is consistent with the progressive transporter loss observed in our study and provides cross-modal validation of our findings. The dramatic mitochondrial and metabolic protein collapse we document has not been a primary focus of these studies. Whether it reflects transcriptional silencing, mitophagy, altered protein stability, or post-translational mechanisms such as succinylation (processes our group has studied in AKI (43–46)) remains an important open question that future proteomics-transcriptomics integration will need to address.

Our global whole-kidney proteomics approach has yielded comprehensive molecular insights; however, an important complementary future direction is the application of spatial proteomics to preserve anatomical context within the kidney’s distinct functional compartments. The Kidney Precision Medicine Project (KPMP) has pioneered integration of multiple spatial technologies to construct a reference kidney tissue atlas, achieving identification of over 2,500 proteins from isolated glomeruli and proximal tubules using near-single-cell proteomics (LCM-microPOTS) and revealing segment-specific proteomic signatures (63, 64). Spatial top-down proteomics approaches combining LCM with nanoliter-scale sample preparation now permit near-cellular spatial resolution over entire tissue sections, identifying proteoforms with differential abundance between compartments (65). Multiplexed imaging platforms including CODEX and spatial MIST have been applied to map protein expression in diabetic kidney disease and fibrotic CKD, enabling spatial proximity analysis of fibrosis-associated cellular markers (66, 67). Future studies integrating our proteomics findings with spatial approaches, including MALDI-MS Imaging (68), which our group has applied to metabolite mapping in kidney tissue (69), would substantially advance mechanistic understanding of the compartment-specific responses driving AKI-to-CKD transition.

The coordinated loss of peroxisomal proteins in both AKI and CKD is of particular interest. Peroxisomes perform the β-oxidation of very-long-chain and branched-chain fatty acids, ether lipid synthesis, and hydrogen peroxide detoxification, and they cooperate closely with mitochondria in proximal tubule lipid metabolism. In our data, the depletion of peroxisomal β-oxidation enzymes, peroxins, and catalase was evident after ischemia-reperfusion and was also present, although to a lesser degree, after ureteral obstruction. This pattern suggests that peroxisomal dysfunction is an early and shared component of kidney injury that may contribute to lipid accumulation, oxidative stress, and the failure of tubular repair. Strategies that restore peroxisomal function, for example through PPARα activation or by targeting individual peroxisomal enzymes, may therefore represent therapeutic opportunities across the AKI-to-CKD continuum.

Our data also reveal a SASP-like signature in both the AKI and CKD models. Senescent tubular cells accumulate after ischemic injury and in fibrotic kidneys, and their secretome promotes inflammation, fibrosis, and maladaptive repair (26) (27) (28) (29). The consistent up-regulation of injury and SASP-associated factors such as Havcr1, Spp1, Igfbp7, Plau, Vcam1, and Icam1, together with lysosomal cathepsins and galectins, indicates that a senescence-associated secretory program is engaged in both acute and chronic injury, and is strongest after ischemia-reperfusion. Although canonical cell-cycle arrest markers were below the detection limit of our tissue proteomics workflow, these findings are in line with the emerging view of senescence as a link between kidney injury, kidney aging, and CKD progression. Importantly, senescent cells can be targeted pharmacologically, and senolytic interventions have shown benefit in preclinical models of kidney aging and disease (30). The SASP-like proteins identified here may serve as tissue readouts to evaluate such interventions.

## Conclusions

This comprehensive proteomic analysis of AKI and CKD mouse models provides several key insights into kidney injury pathophysiology with direct implications for diagnosis and therapy.

### Conserved core kidney injury response

We demonstrate that nearly half (46%) of altered proteins are shared between AKI and CKD, revealing a conserved core injury response that operates independently of the initial insult mechanism. Most striking is the universal metabolic dysfunction signature: both acute ischemic injury and chronic obstructive injury result in profound down-regulation of mitochondrial proteins, cellular respiration machinery, fatty acid oxidation enzymes, and solute transporters. This convergent metabolic collapse likely reflects the kidney’s unique vulnerability due to its high energy demands, and suggests that mitochondrial protection strategies may benefit patients with kidney injury regardless of etiology.

### Disease-specific signatures distinguish acute from chronic injury

Despite the substantial overlap, our analysis reveals important disease-specific features. AKI is characterized by acute stress signatures including chromatin reorganization, DNA damage response activation, and acute inflammation, reflecting the immediate cellular response to ischemia-reperfusion. In contrast, CKD shows signatures of progressive tissue damage including sustained fibrosis, complement cascade activation, and coagulation pathway engagement. These distinct molecular profiles provide a framework for understanding why some patients recover from acute injury while others progress to chronic disease.

### Novel progression, phase-specific, and peroxisomal markers

A major contribution of this work is the identification of 95 candidate progression markers showing greater increases in CKD than in AKI, and 14 opposite-direction proteins with divergent patterns between disease phases. Additionally, subcellular localization analysis identified a coordinated loss of peroxisomal proteins, including the β-oxidation enzymes Acox1 and Ehhadh, in both AKI and CKD, alongside a broader collapse of proximal tubule metabolic enzymes spanning mitochondrial and peroxisomal compartments, collectively representing a novel metabolic signature mechanistically linked to tubular lipotoxicity and fibrotic progression. The progression markers, phase-specific proteins, and metabolic signatures collectively provide candidate biomarkers for disease monitoring, phase discrimination, and therapeutic targeting.

### Senescence-associated secretory signature

Both AKI and CKD displayed a SASP-like signature, including injury and SASP-associated factors, lysosomal cathepsins, and galectins, which was most pronounced after ischemia-reperfusion. This finding connects kidney injury to cellular senescence, a hallmark of aging, and suggests that senolytic or senomorphic interventions may help to improve kidney health after acute injury and to slow progression to CKD.

### Future directions

Several avenues for future investigation emerge from this work. Validation of the identified progression and phase-specific markers in human kidney biopsy samples and clinical cohorts will be essential for translation. Temporal proteomics studies capturing the dynamics of protein expression during injury, repair, and progression would complement our single-time-point analysis. The peroxisomal dysfunction signature warrants functional investigation, in particular, whether restoration of peroxisomal β-oxidation (e.g., via PPARα activation) can mitigate tubular lipotoxicity and attenuate AKI-to-CKD transition. Whether senolytic interventions can reduce the SASP-like signature and improve kidney outcomes after AKI also warrants investigation. Cell-type-specific and spatial proteomic approaches could resolve which kidney cell populations drive the observed signatures, and functional validation of candidate markers such as Mfn2, Brat1, and Acox1 as therapeutic targets could open new avenues for preventing CKD progression.

## Summary

In conclusion, our comparative proteomic analysis of AKI and CKD reveals a shared core injury response dominated by metabolic dysfunction alongside disease-specific signatures that distinguish acute from chronic injury. The identification of candidate progression markers, opposite-direction proteins, a coordinated loss of peroxisomal proteins in both AKI and CKD, and a SASP-like senescence signature advances our molecular understanding of kidney injury pathophysiology and provides a rich resource for biomarker discovery and therapeutic target identification in AKI and CKD, including strategies to restore peroxisomal function and senolytic approaches to improve kidney health.

## Materials and Methods

### Sex as a biological variable

Our study exclusively examined male mice. We currently cannot assess sex-specific differences in AKI mechanisms and transition to CKD, although females seem more protected from CKD progression (70, 71). Sex may influence the molecular mechanisms of the disease continuum. Female mice should therefore also be investigated to determine the relevance of these findings.

### Mouse kidney tissues

Wild-type mice were obtained from the Jackson Laboratory (B6129SF1/J, stock no. 101043 for renal IRI; 129S1/SvImJ). Age-matched 9-to 14-week-old male mice were used throughout the study. The University of Pittsburgh Institutional Animal Care and Use Committee approved all experiments (approval no. 22112009).

### Ischemia-reperfusion injury (IRI) AKI Model

Ischemic AKI was induced by a unilateral renal ischemia-reperfusion injury (IRI) with a delayed contralateral nephrectomy as previously described (44, 46). Briefly, mice were anesthetized with 2% isoflurane. Core body temperature was maintained at 36–37°C throughout with a water-heating circulation pump system (EZ-7150; BrainTree Scientific) and an infrared heat lamp. Ethiqa® extended-release buprenorphine 1.3 mg/mL was administered subcutaneously for analgesia (3.25 mg/kg). With aseptic technique, a dorsal incision was made to expose the kidney, and renal ischemia for 22 minutes was induced by clamping of the left kidney pedicle with a nontraumatic microvascular clamp (18055–04; Fine Science Tools). Renal reperfusion was visually verified. Contralateral nephrectomy was performed at day 6. Mice were euthanized at day 7 to harvest the IRI kidney. Serum creatinine, BUN, and other biochemical markers were analyzed by the Kansas State Veterinary Diagnostic Laboratory.

### Unilateral ureteral obstruction (UUO) CKD model

To induce renal fibrosis, mice were subjected to UUO. Following anesthesia with 2% isoflurane and analgesia as above, a midline laparotomy was performed, the left ureter was located by blunt dissection, and the ureter was ligated. Animals recovered for 4 days prior to sacrifice. Non-obstructed contralateral kidneys served as paired controls. UUO represents an accelerated fibrosis model that captures key molecular features of CKD-associated fibrotic remodeling, though it does not recapitulate the prolonged timeline of human CKD progression (8).

### Proteomic sample preparation – Acute kidney injury cohort

Kidney tissues from mice subjected to a 22-min ischemia-reperfusion (n = 4) and control mice (n = 3) were collected. Frozen kidneys were immersed in 500 μL of lysis buffer containing 8 M urea, 200 mM triethylammonium bicarbonate (TEAB), pH 8, 75 mM sodium chloride, 1 μM trichostatin A, 3 mM nicotinamide, and 1x protease/phosphatase inhibitor cocktail (Thermo Fisher Scientific, Waltham, MA), and homogenized for 2 cycles with a Bead Beater TissueLyser II (Qiagen, Germantown, MD) at 25 Hz for 4 min each. Lysates were clarified by spinning at 15,700 x *g* for 15 min at 4°C, and the supernatant containing the soluble proteins was collected. After protein concentration determination using BCA kit (Thermo Fisher Scientific, Waltham, MA), 200 μg protein samples were brought to the same volume using water. Samples were solubilized using 4% SDS, 50 mM TEAB. Proteins were reduced using 20 mM dithiothreitol (DTT; 10 min at 50°C followed by 10 min at RT) and alkylated using 40 mM iodoacetamide (IAA; 30 min at RT in the dark). Samples were acidified with a final concentration of 1.2% phosphoric acid and diluted with seven volumes of S-trap buffer (90% methanol in 100 mM TEAB, pH ∼7). Samples were then loaded onto the S-trap mini spin columns (Protifi, Farmingdale, NY) and spun at 4,000 x *g* for 10 seconds. The S-trap columns were washed with S-trap buffer twice at 4,000 x *g* for 10 seconds each, before adding a solution of sequencing grade trypsin (Promega, San Luis Obispo, CA) in 50 mM TEAB at a 1:25 (w/w) enzyme:protein ratio. After a 1-hour incubation at 47°C, trypsin solution was added again at the same ratio, and proteins were digested overnight at 37°C. Peptides were sequentially eluted with 50 mM TEAB (spun through for 1 min at 1,000 x *g*), 0.5% formic acid (FA) in water (spun through for 1 min at 1,000 x *g*), and 50% acetonitrile (ACN) in 0.5% FA (spun through for 1 min at 4,000 x *g*). After vacuum drying, samples were resuspended in 0.2% FA in water and desalted with Oasis 10 mg Sorbent Cartridges (Waters, Milford, MA). They were vacuum dried and resuspended in 0.2% FA in water at a final concentration of 1 μg/μL. Finally, indexed retention time standard peptides (iRT; Biognosys, Schlieren, Switzerland) were spiked in the samples according to the manufacturer’s instructions (72).

### Proteomic sample preparation – Chronic kidney disease cohort

Both unilateral ureteral obstructed kidney and contralateral control kidney tissues from wild-type mice (n=4) were collected. Frozen kidneys were immersed in 700 μL of lysis buffer and homogenized as described above. Lysates were clarified by spinning at 15,700 x *g* for 15 min at 4°C, and the supernatant containing the soluble proteins was collected. After protein concentration determination using BCA (Thermo Fisher Scientific), 3.1 mg protein of each sample were reduced using 20 mM DTT in 50 mM TEAB for 30 min at 37 °C, and after cooling to room temperature, alkylated with 40 mM IAA for 30 min at room temperature in the dark. Samples were diluted 10-fold with 50 mM TEAB and proteins were digested overnight with a solution of sequencing-grade trypsin (Promega, San Luis Obispo, CA) in 50 mM TEAB at a 1:50 (wt:wt) enzyme:protein ratio at 37°C. This reaction was quenched with 1% formic acid (FA) and the sample was clarified by centrifugation at 1,800 x *g* for 15 min at room temperature. Clarified peptide samples were desalted with Oasis 30 mg Sorbent Cartridges (Waters, Milford, MA). Aliquots of 100 μg of each peptide elution were vacuum dried, desalted again with Oasis 10 mg Sorbent Cartridges (Waters), vacuum dried, and finally re-suspended in 0.2% FA at a final concentration of 1 μg/μL. Indexed retention time standard peptides (iRT; Biognosys) were spiked in the samples according to the manufacturer’s instructions (72).

### DIA-MS analysis

LC-MS/MS analyses were performed on a Dionex UltiMate 3000 system coupled to an Orbitrap Eclipse Tribrid mass spectrometer (both from Thermo Fisher Scientific, San Jose, CA). The solvent system consisted of 2% ACN, 0.1% FA in water (solvent A) and 98% ACN, 0.1% FA in water (solvent B). Peptides (200 ng) were loaded onto an Acclaim PepMap 100 C18 trap column (75 µm x 20 mm, 3 µm particle size; Thermo Fisher Scientific) over 10 min at 2 µL/min with 100% solvent A. Peptides were eluted on an Acclaim PepMap 100 C18 analytical column (75 µm x 50 cm, 3 µm particle size; Thermo Fisher Scientific) at 0.3 µL/min using the following gradient of solvent B: 2% for 10 min, linear from 2% to 20% in 125 min, linear from 20% to 32% in 40 min, up to 80% in 1 min, 80% for 9 min, and down to 2% in 1 min. The column was equilibrated with 2% of solvent B for 29 min, with a total gradient length of 215 min. All samples were analyzed in data-independent acquisition (DIA) mode (73–75). Full MS spectra were collected at 120,000 resolution (AGC target: 3e6 ions, maximum injection time: 60 ms, 350-1,650 m/z), and MS2 spectra at 30,000 resolution (AGC target: 3e6 ions, maximum injection time: Auto, NCE: 27, fixed first mass 200 m/z). The isolation scheme consisted of 26 variable windows covering the 350-1,650 m/z range with an overlap of 1 m/z (DIA isolation scheme) (73).

### Data analysis with directDIA (Spectronaut)

Data were processed in Spectronaut (version 19.4.241104.62635) using directDIA against the *Mus musculus* reference proteome (UniProtKB-TrEMBL, 54,910 entries; accessed 05/18/2024) (76). Trypsin/P was set as digestion enzyme with two missed cleavages. Cysteine carbamidomethylation was set as a fixed modification; methionine oxidation and protein N-terminus acetylation were variable modifications. Data extraction parameters were set as dynamic. Identification was performed at 1% precursor and protein q-value.

Quantification was based on extracted ion chromatograms (XICs) of 3–6 MS2 fragment ions with local normalization and q-value sparse filtering. Protein groups were required to have at least two unique peptides. Differential analysis is described in the Statistics section **(Supplemental Table 1)**.

### Clustering analysis and heatmap generation

Principal component analysis (PCA) was performed using the stats R package. The heatmap was generated on all quantified protein groups using the heatmap.2() function from the gplots R package.

### Over-representation analysis

An over-representation analysis (ORA) was performed using ConsensusPathDB (48, 49) to determine which gene ontology (GO) terms were significantly enriched. The background proteome was defined as all quantified protein groups in this dataset. GO terms were considered significant as described in the Statistics section. Dot plots were generated using the ggplot2 R package (77).

### LOPIT organellar protein distribution

The hyperLOPIT2015 pluripotent stem cell dataset was downloaded in RStudio using the pRolocData R package (Bioconductor) and plotted with the pRoloc R package (78). The LOPIT reference dataset was generated from mouse pluripotent stem cell lysate fractionated by density gradient ultracentrifugation, with proteins assigned to organelles based on similarities in distribution to well-annotated organelle marker proteins (54, 79, 80). The t-SNE algorithm was used to reduce the multi-dimensional LOPIT proteomic data to a 2D map where proteins cluster by subcellular similarity (53, 81). All LOPIT protein assignments were used as originally determined and without additional refinement.

### Weighted gene co-expression network analysis (WGCNA)

Protein co-expression network analysis was performed using protein abundance with the WGCNA R package (52) as described by Gu et al. (82). A soft power of 14, minimum cluster size of 30, and no module merging were applied. Module eigengenes were correlated with experimental condition traits using Pearson correlation, and significance was assessed by Student’s asymptotic p-values. AKI- and CKD-associated modules were defined by r > 0.8 and p < 0.05 **(Supplemental Tables 5 and 6)**. Cluster proteins were submitted to ORA as described above.

### Peroxisomal and senescence protein panels

Peroxisomal proteins were defined as quantified protein groups with established peroxisomal localization, grouped by function (β-oxidation, carnitine and acyl-CoA metabolism, α-oxidation, oxidases and antioxidant defense, ether lipid synthesis, and peroxisome biogenesis, import, and membrane transport); proteins with predominant localization to other compartments were excluded **(Supplemental Table 7)**. Senescence and SASP-associated proteins were compiled from the SASP Atlas (58) and the SenMayo gene set (59), together with canonical senescence markers, and grouped by category **(Supplemental Table 8)**. For both panels, differential abundance values were taken from the differential analysis of all quantified protein groups **(Supplemental Table 1)**, using the same significance criteria (q-value < 0.01 and absolute log2(fold-change) > 0.58).

### Use of AI

Claude (Anthropic; Claude Opus 4.5 and Claude Opus 5.5, model identifier claude-opus-5-5) was used between February 2026 and November 2026 to assist with figure layout, synthesis and visualization of pathway enrichment results, compilation of the curated peroxisomal and senescence protein panels from the supplemental data tables, and editing of the manuscript text. All underlying statistical analyses were performed by the authors as described above. The authors reviewed and verified all AI-assisted output and take full responsibility for the content of this manuscript.

### Statistics

Differential analysis was performed using an unpaired t-test, and p-values were corrected for multiple testing using the Storey method (83, 84). Significance was defined as q-value < 0.01 and absolute log2(fold-change) > 0.58. Pearson correlation analysis was performed on the log2(fold-change) of all protein groups significantly altered in both AKI (AKI-IRI vs Sham-Nephrectomy) and CKD (CKD-Obstructed vs Non-Obstructed) in R using the cor.test() function. For the ORA, p-values were determined by a hypergeometric test and corrected for multiple testing using the false discovery rate approach; GO terms with q ≤ 0.001 and term_level = 4, or p ≤ 0.01 and count ≥ 5, were considered significant. For the WGCNA, module eigengenes were correlated with experimental condition traits using Pearson correlation, and significance was assessed by Student’s asymptotic p-values; trait-associated modules were defined by r > 0.8 and p < 0.05. For each module, eigengene values were compared across the four groups using the non-parametric Kruskal-Wallis test, with p < 0.05 considered significant.

### Study approval

All animal experiments were approved by the IACUC protocol with approval number 22112009.

## Data Availability

The mass spectrometry data that support the findings of this study are available via the MassIVE repository (UCSD) at https://massive.ucsd.edu/ProteoSAFe/dataset.jsp?task=25bb7fe9e595417089ae9a51879b5b2d or via the FTP link ftp://MSV000101752@massive-ftp.ucsd.edu (MassIVE ID: MSV000101752; ProteomeXchange ID: PXD078127). [Note to reviewers: Username: MSV000101752_reviewer; Password: winter].

## Acknowledgments

This work was supported by NIH/NIDDK R01 DK121758, R01 DK134346 and by R25 DK119180 (PI: S.S.L.) and by the NIH/NCRR shared instrumentation grant S10 OD028654 (B.S.).

## Supplemental Figure Legends

**Supplemental Figure 1. Overview of shared and disease-specific proteome signatures.** (A) Proteome coverage and overlap: 5,985 protein groups were quantified, 3,194 were significantly altered in AKI and/or CKD, and 1,475 (46%) were shared between AKI and CKD. (B) Disease-specific signatures: 1,254 AKI-specific (836 up-regulated and 418 down-regulated) and 465 CKD-specific (242 up-regulated and 223 down-regulated) protein groups, with representative biological functions. (C) Breakdown of the 1,475 shared proteins into 622 up-regulated, 839 down-regulated, and 14 opposite-direction protein groups, with representative biological processes. (D) Overview of key findings. Statistical criteria: q-value < 0.01 and |log2(fold-change)| > 0.58.

**Supplemental Figure 2. Slope plots of candidate AKI-to-CKD progression markers.** (A) Candidate progression markers that were up-regulated in both AKI and CKD, with a fold-change at least 1.5-fold higher in CKD (n=95; selected proteins shown). (B) Proteins with progressive loss that were down-regulated in both AKI and CKD, with a fold-change at least 1.5-fold lower in CKD (n=93; selected proteins shown). Open circles, AKI log2(fold-change); filled circles, CKD log2(fold-change). Proteins are grouped by biological function. Statistical criteria: q-value < 0.01 and |log2(fold-change)| > 0.58.

**Supplemental Figure 3. Weighted gene co-expression network analysis (WGCNA) module-trait relationships and module eigengenes.** (A) Module-trait relationships for the modules associated with AKI and CKD. Values are Pearson correlation coefficients, with Student asymptotic p-values in parentheses. Tier 1 modules (r > 0.8 and p < 0.05; dark arrows) were Dark red, Light cyan, and Royal blue for AKI, and Red, Tan, Black, and Cyan for CKD; tier 2 modules (r > 0 and p < 0.05) are indicated by light arrows. Soft power = 14; minimum cluster size = 30; no module merging. (B) Module eigengene values of the three AKI-associated and four CKD-associated modules for each replicate in the four experimental groups (AKI-IRI, Sham-Nephrectomy, CKD-Obstructed, and Non-Obstructed), with Kruskal-Wallis p-values. The representative GO term of each module, determined by over-representation analysis, is indicated (p ≤ 0.01 and count ≥ 5).

**Supplemental Figure 4. LOPIT reference map used for subcellular localization analysis.** Mouse pluripotent stem cell LOPIT reference map (t-SNE dimensionality reduction) (54), showing the nucleus (chromatin and non-chromatin), mitochondria, endoplasmic reticulum/Golgi apparatus, plasma membrane, cytosol, 40S and 60S ribosomes, peroxisome, lysosome, endosome, extracellular matrix, actin cytoskeleton, and proteasome compartments.

