## Supplemental Figure 1 for "Proteomic Evaluation of the Progression from Acute Kidney Injury to Chronic Kidney Disease"

### Disease-specific and shared proteome signatures

#### A Proteome coverage and overlap

**5,985**
Protein groups quantified

**3,194**
Significantly altered (AKI and/or CKD)

**1,475 (46%)**
Shared between AKI and CKD

#### B Disease-specific signatures

**AKI-specific: 1,254 proteins**

**Up-regulated (836)**
- Chromatin organization
- DNA damage response
- ECM organization and innate immunity
- Metabolic reprogramming

**Down-regulated (418)**
- Amino acid metabolism
- Gluconeogenesis
- Solute carriers
- Peroxisomal proteins

**CKD-specific: 465 proteins**

**Up-regulated (242)**
- Fibrosis and ECM
- Complement cascade
- Inflammation

**Down-regulated (223)**
- Solute transporters
- Mitochondrial proteins
- Tubular markers

#### C Breakdown of the 1,475 shared proteins

**Shared up-regulated: 622**
- ECM organization
- Adaptive immune response
- Cytokine production
- DNA-binding transcription factor activity
- Circulatory system development
- Cell migration

**Shared down-regulated: 839**
- Branched-chain amino acid metabolism
- Respiratory electron transport chain
- Cellular respiration and fatty acid oxidation
- Acyl-CoA metabolism
- Mitochondrion organization
- Transmembrane and ion transport

**14 shared proteins changed in opposite directions (6 up in AKI and down in CKD; 8 the reverse)**

#### D Key findings

1. A conserved core injury response: 46% of altered proteins are shared between AKI and CKD.
2. Metabolic collapse in both models: mitochondrial and peroxisomal proteins down-regulated (peroxisomal: 45 in AKI and 28 in CKD, none up-regulated).
3. AKI shows acute stress signatures: chromatin remodeling, DNA damage response, and acute inflammation.
4. CKD shows chronic remodeling: sustained fibrosis, complement activation, and coagulation.
5. A SASP-like senescence signature is shared by both models and is stronger in AKI.

Statistical criteria: $q$-value $< 0.01$ and $|\log_2(FC)| > 0.58$; GO enrichment: ConsensusPathDB, $q$-value $< 0.001$

**Supplemental Figure 1. Overview of shared and disease-specific proteome signatures.** (A) Proteome coverage and overlap: 5,985 protein groups were quantified, 3,194 were significantly altered in AKI and/or CKD, and 1,475 (46%) were shared between AKI and CKD. (B) Disease-specific signatures: 1,254 AKI-specific (836 up-regulated and 418 down-regulated) and 465 CKD-specific (242 up-regulated and 223 down-regulated) protein groups, with representative biological functions. (C) Breakdown of the 1,475 shared proteins into 622 up-regulated, 839 down-regulated, and 14 opposite-direction protein groups, with representative biological processes. (D) Overview of key findings. Statistical criteria: q-value $< 0.01$ and $|\log2(\text{fold-change})| > 0.58$.
