## Supplemental Figure 2 for "Proteomic Evaluation of the Progression from Acute Kidney Injury to Chronic Kidney Disease"

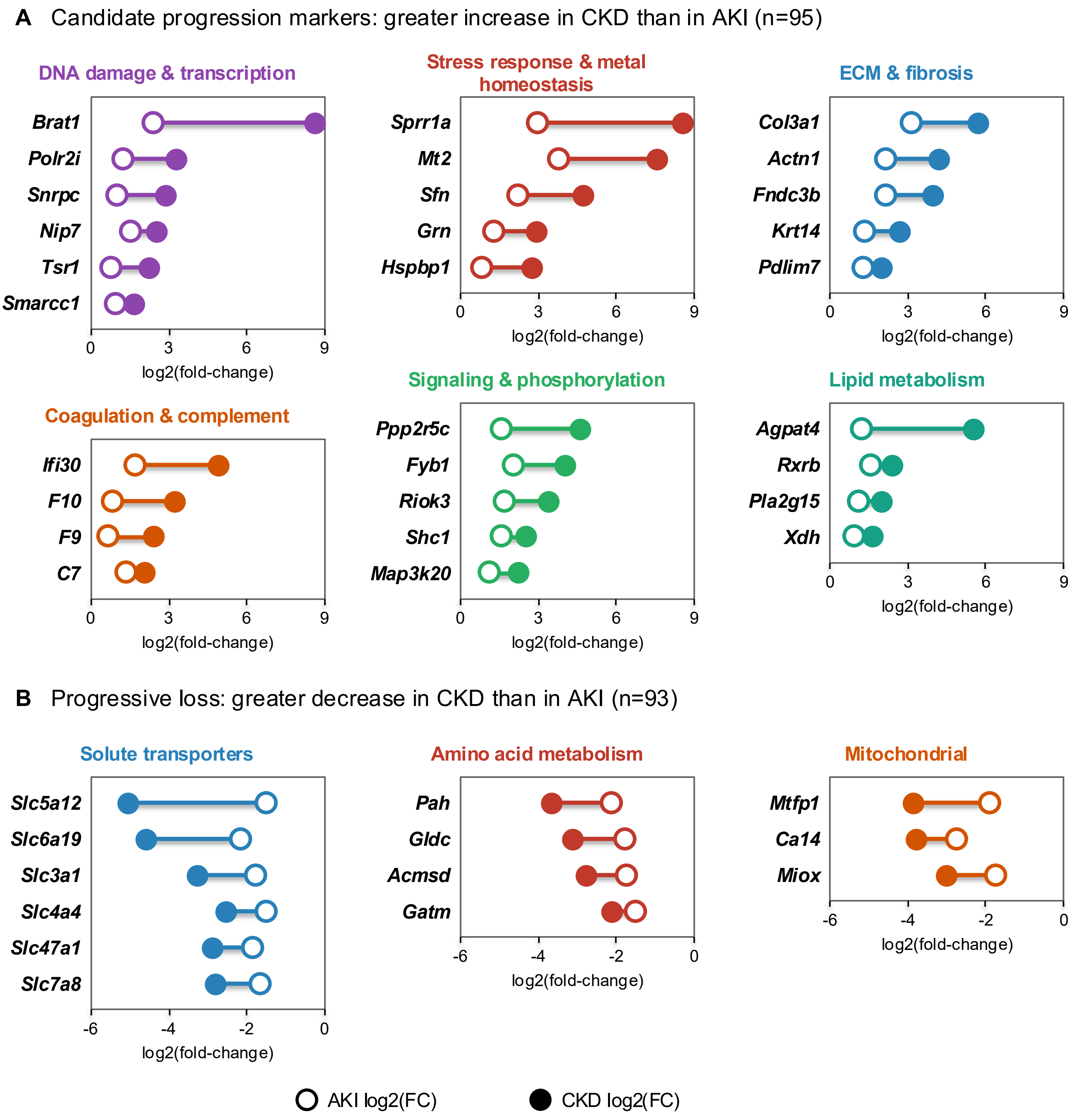


Selected proteins shown; q-value < 0.01 and |log2(FC)| > 0.58 in both comparisons, with at least 1.5-fold difference between CKD and AKI.

**Supplemental Figure 2. Slope plots of candidate AKI-to-CKD progression markers.** (A) Candidate progression markers that were up-regulated in both AKI and CKD, with a fold-change at least 1.5-fold higher in CKD (n=95; selected proteins shown). (B) Proteins with progressive loss that were down-regulated in both AKI and CKD, with a fold-change at least 1.5-fold lower in CKD (n=93; selected proteins shown). Open circles, AKI log2(fold-change); filled circles, CKD log2(fold-change). Proteins are grouped by biological function. Statistical criteria: q-value < 0.01 and |log2(fold-change)| > 0.58.
