## Supplemental Figure 3 for "Proteomic Evaluation of the Progression from Acute Kidney Injury to Chronic Kidney Disease"

**A**

### WGCNA of AKI and CKD Proteomes

### Module-trait relationships

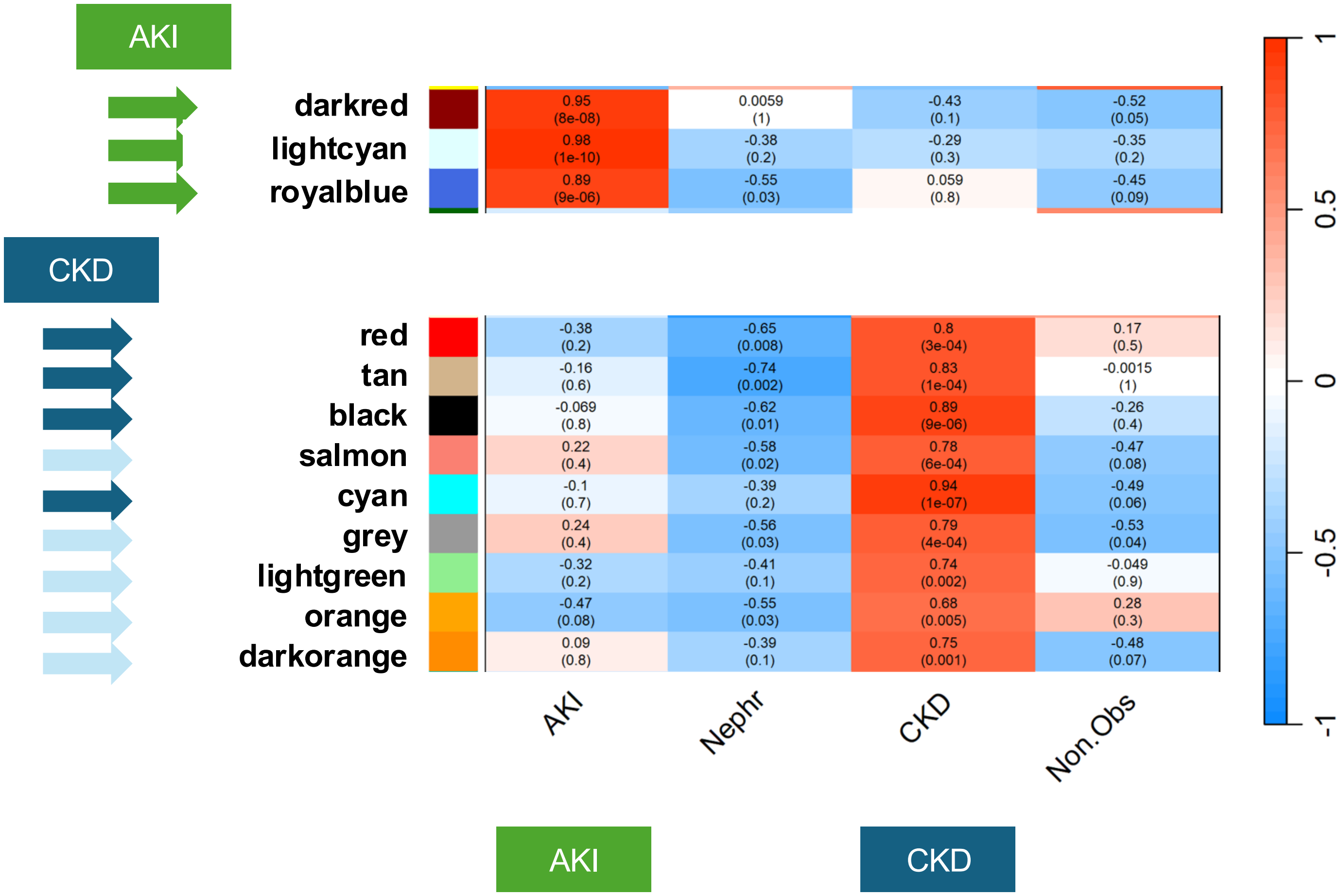


<u>Values in heatmap:</u>
Pearson's correlation coefficient
Student asymptotic p-value

**3 modules** associated with **AKI**
**4 modules** associated with **CKD**

<u>**Tier 1:** r > 0.8 and p < 0.05</u>
**Tier 2:** r > 0 and p < 0.05

Soft power = 14; Min cluster size = 30; No merging

Langfelder P. and Horvath S., *BMC Bioinformatics*, 2008

**Supplemental Figure 3 (continued)**

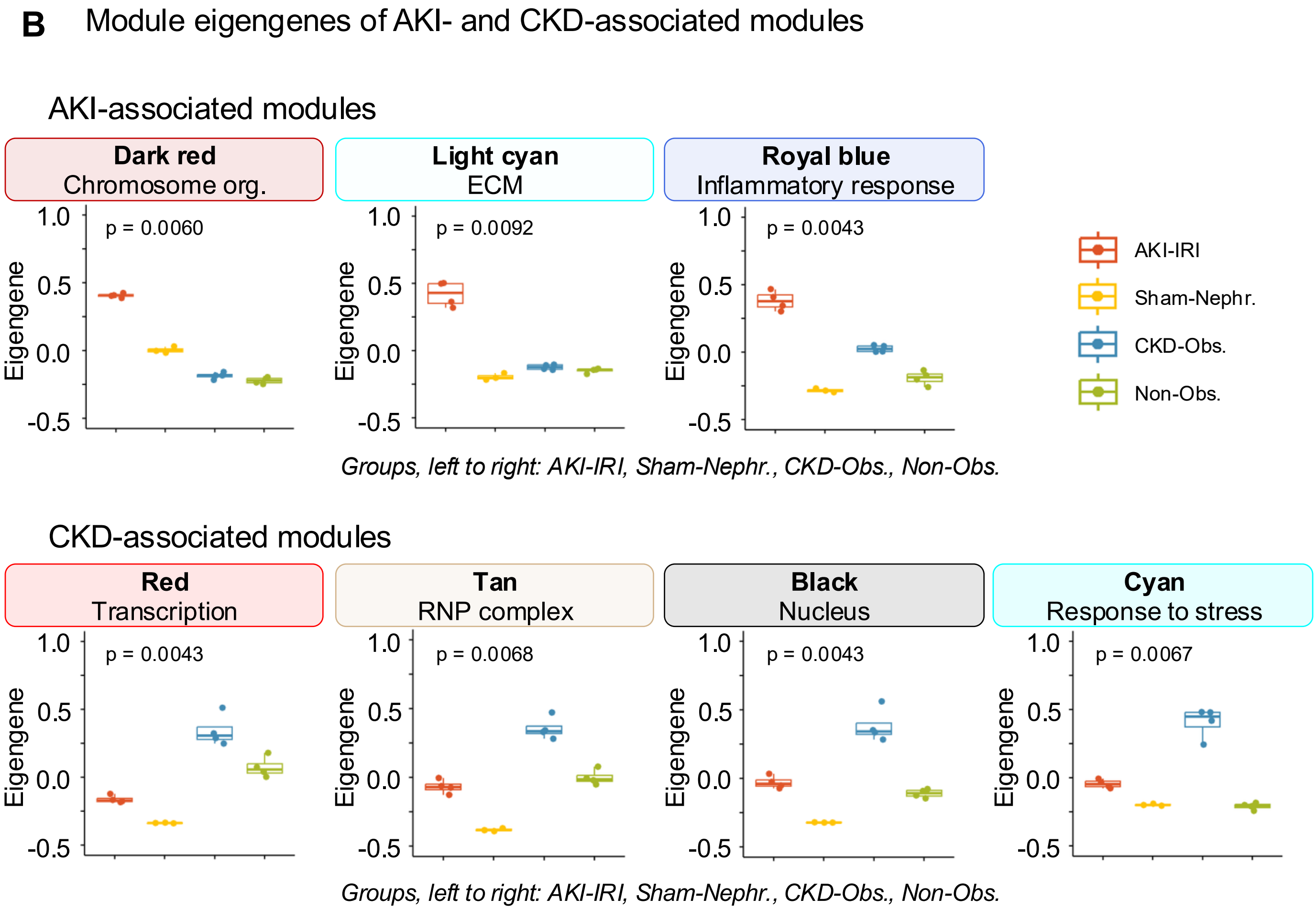


**Supplemental Figure 3. Weighted gene co-expression network analysis (WGCNA) module-trait relationships and module eigengenes.** (A) Module-trait relationships for the modules associated with AKI and CKD. Values are Pearson correlation coefficients, with Student asymptotic p-values in parentheses. Tier 1 modules ($r > 0.8$ and $p < 0.05$; dark arrows) were Dark red, Light cyan, and Royal blue for AKI, and Red, Tan, Black, and Cyan for CKD; tier 2 modules ($r > 0$ and $p < 0.05$) are indicated by light arrows. Soft power = 14; minimum cluster size = 30; no module merging. (B) Module eigengene values of the three AKI-associated and four CKD-associated modules for each replicate in the four experimental groups (AKI-IRI, Sham-Nephrectomy, CKD-Obstructed, and Non-Obstructed), with Kruskal-Wallis p-values. The representative GO term of each module, determined by over-representation analysis, is indicated ($p \leq 0.01$ and count $\geq 5$).
