## Supplemental Figure 4 for "Proteomic Evaluation of the Progression from Acute Kidney Injury to Chronic Kidney Disease"

### Subcellular Localization of Quantified Proteins

#### LOPIT Map: Reference for Mouse

*Localization of Organelle Proteins by Isotopic Tagging*

Christoforou *et al.*, *Nature Communications*, 2016
Burton *et al.*, *Proteomes*. 2022

##### Mouse Pluripotent Stem Cell LOPIT Map

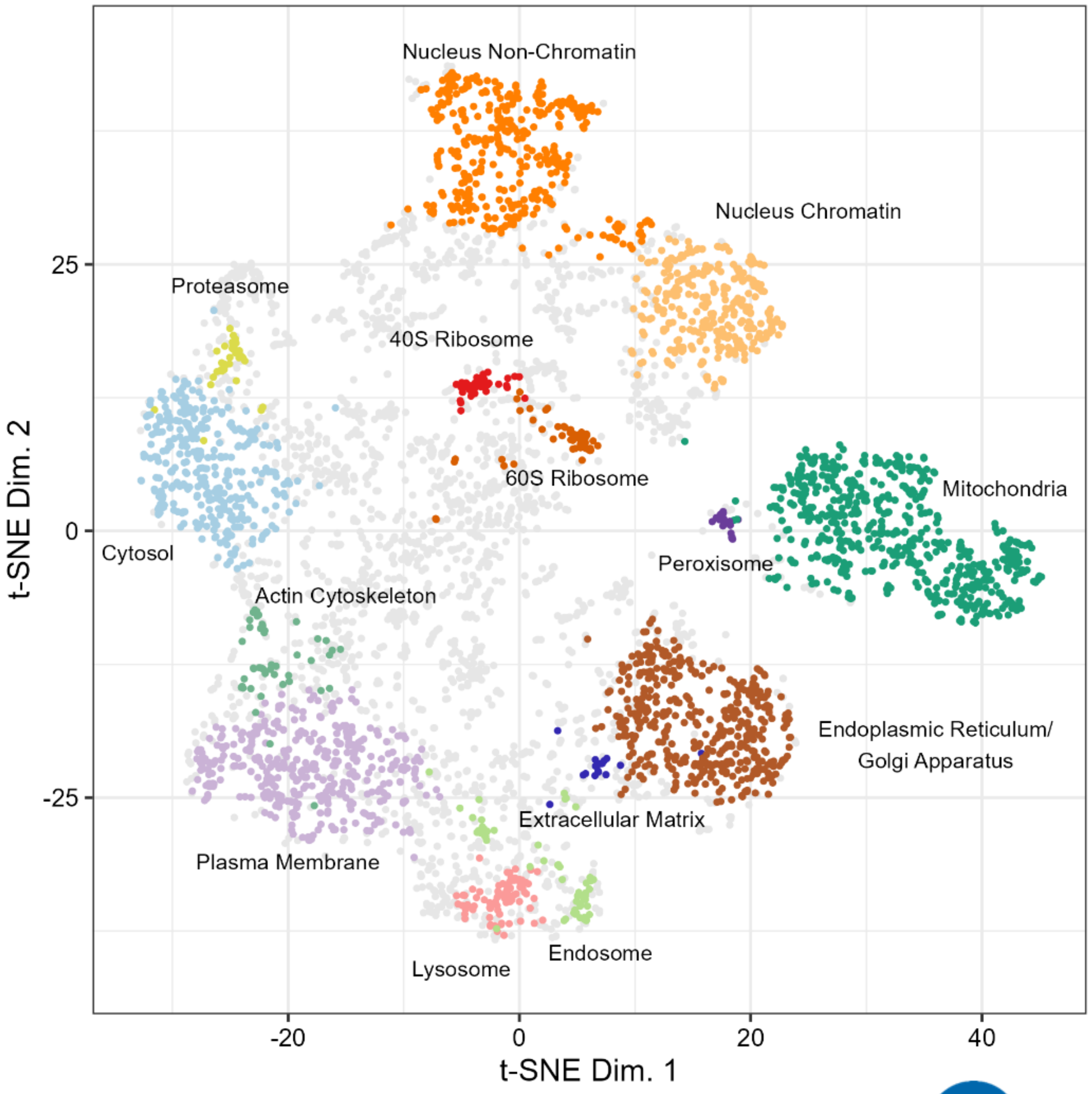


Buck

**Supplemental Figure 4. LOPIT reference map used for subcellular localization analysis.** Mouse pluripotent stem cell LOPIT reference map (t-SNE dimensionality reduction), showing the nucleus (chromatin and non-chromatin), mitochondria, endoplasmic reticulum/Golgi apparatus, plasma membrane, cytosol, 40S and 60S ribosomes, peroxisome, lysosome, endosome, extracellular matrix, actin cytoskeleton, and proteasome compartments (Christoforou et al., Nat Commun 2016).
